# Targeted suppression of SPP1 inhibits tumor invasion and metastasis in NRF2 hyperactivated cisplatin resistant HNSCC

**DOI:** 10.1101/2025.05.28.656679

**Authors:** Mutsuki Kawabe, Sujuan Yang, Lorena Gomez Bolanos, Shiro Takamatsu, Sylvia Flores, Patricia D. Castro, Tsung-You Tsai, Arisa Nishikawa-Kaga, Mitchell J. Frederick, Vlad Sandulache, Humam Kadara, Jeffrey N. Myers, Abdullah A. Osman

**Affiliations:** Department of Head and Neck Surgery, The University of Texas MD Anderson Cancer Center, Houston, Texas; Department of Otolaryngology-Head and Neck Surgery, Baylor College of Medicine, Houston, Texas; Department of Translational Molecular Pathology, The University of Texas MD Anderson Cancer Center, Houston, Texas; Department of Gynecologic Oncology and Reproductive Medicine, The University of Texas MD Anderson Cancer Center, Houston, Texas

**Keywords:** Cisplatin, CDDP, Head and neck cancer, Osteopontin, SPP1, KEAP1, NRF2, integrins, CD44

## Abstract

**Purpose:** Cisplatin is the gold standard systemic agent for definitive treatment of HNSCC. The purpose of this study was to investigate the role of SPP1 in the progression and metastasis of cisplatin-resistant HNSCC, particularly in the context of NRF2 hyperactivation.

**Experimental Design:** CDDP resistant HNSCC cell lines stably expressing various shRNA SPP1 constructs were generated. Clonogenic survival assays and a mouse model of oral cancer were used to examine the impact of silenced SPP1 *in vitro* and *in vivo* on CDDP sensitivity and tumor progression. Western blotting, cell invasion, functional proteomics RPPA and spatial transcriptomic analyses were performed to identify molecular interactions and metastatic signaling pathways.

**Results:** Targeted suppression of SPP1 improved cisplatin sensitivity, inhibited tumor invasion and metastasis *in vitro* and *in vivo* as well as several metastatic signaling proteins in NRF2- hyperactivated HNSCC. Spatial transcriptomic analysis revealed a potential mechanistic interaction between SPP1, integrins and CD44 receptors in primary and metastatic HNSCC. Spatial annotation and enrichment analyses using HALLMARK revealed gene set signatures of interferon and EMT present in cell clusters with SPP1 expression in both the primary tumor and lung metastases. Finally, increased expression of SPP1 was found to be poor prognostic factor and significantly correlated with *NFE2L2/KEAP1* mutational status and higher tumor grade in HNSCC patients.

**Conclusions:** Targeting dysregulated SPP1 improved cisplatin sensitivity, inhibited tumor invasion and metastasis in NRF2-hyperactivated HNSCC. These findings highlight the therapeutic potential of targeting SPP1 and the need for developing specific SPP1 inhibitors to improve outcomes for patients with HNSCC.

**Translational Relevance:** Cisplatin-based chemotherapy remains the standard of care for patients with locally advanced head and neck squamous cell carcinoma. Cisplatin resistance often leads to treatment failure and metastasis which accounts for the majority of mortality associated with this disease. Unfortunately, there are no effective therapeutic agents to overcome cisplatin resistance in HNSCC. Here, we present compelling evidence that targeting SPP1 (also known as osteopontin) inhibits cisplatin resistance, tumor progression and metastasis in NRF2-hyperactivated HNSCC, positioning it as a potential therapeutic target to improve treatment outcomes in HNSCC.

## Introduction

Head and neck squamous cell carcinoma (HNSCC) is the seventh leading cause of cancer deaths worldwide, with 700,000 cases diagnosed per year (1,2), and the overall 5-year survival remains ∼50% despite efforts to improve therapeutic regimens (3). In the recurrent/metastatic setting, HNSCC is nearly universally fatal. Although immune checkpoint inhibitors have been approved for recurrent metastatic HNSCC, only a small subset of patients (<20%) benefit from this treatment (4). Cisplatin (CDDP) is the gold standard systemic agent for definitive treatment of HNSCC and when combined with immune checkpoint inhibitors (ICIs) for recurrent/metastatic tumors in patients with low combined positive score (CPS>1) of PD-L1 expression in their tumors (5). Unfortunately, continuous exposure to cisplatin is frequently associated with substantial toxicity and can result in the development of acquired resistance (both extrinsic and intrinsic), and therefore some HNSCC tumors show lack of response to cisplatin (6–8). This phenotype has recently been shown by us and others to be driven by hyperactivation of the KEAP1/NRF2 pathway caused by inactivating mutations of *KEAP1* and epigenetic reprogramming of *NFE2L2* (NRF2), resulting in accelerated rates of cervical and lung metastasis *in vivo* in HNSCC preclinical models (7). This is further supported by data suggesting that cells developing cisplatin resistance often exhibit a shift towards a more cellular reductive state (8).

The secreted phosphoprotein 1 (SPP1), also called osteopontin (OPN), is a primary NRF2 target gene overexpressed in a variety of cancer tissues and has been associated with poor prognosis and chemoresistance in several tumor types including lung, breast, and colorectal cancers (9–11). Despite high expression in cancer cells and stroma, there is little evidence supporting mechanistic role of SPP1 in tumor progression and metastasis. SPP1 functions as an extracellular matrix soluble oncoprotein through interaction with integrins and CD44 variants and is also involved in cellular migration, invasion, immune evasion, and tumor metastasis (9,12,13). Our recently published data identified SPP1 as one of the top ten NRF2 downstream target genes that was significantly upregulated *in vivo* in the primary and metastatic tumors in the cisplatin resistant HNSCC cisplatin–treated group compared to the cisplatin-sensitive parental group (7). However, the role of dysregulated SPP1 in cisplatin-resistant HNSCC and development of distant metastasis has not been fully characterized. The current study aims to investigate whether dysregulated SPP1 promotes tumor progression and metastasis, particularly in the context of Nrf2 hyperactivated cisplatin resistant HNSCC. We leveraged our established head and neck cancer preclinical models, which demonstrated *in vitro* and *in vivo* cisplatin resistance that is functionally dependent on dysregulated Nrf2 signaling pathway and associated with high rates of cervical and distant metastasis (7,8,14). In this study we demonstrate that suppressing SPP1 enhances cisplatin sensitivity, inhibits tumor invasion, and reduces metastasis in preclinical models of HNSCC. This effect is linked to ferroptosis-induced cell death rather than apoptosis. Proteomic analysis reveals dysregulation of key oncogenic and metastatic pathways, including MAPK, AKT/mTOR, FAK, and PAK1, in SPP1- silenced cells. Spatial transcriptomics reveals a potential regulatory axis involving tumor-derived SPP1, integrins, and CD44, which is validated by co-immunoprecipitation analysis. TCGA RNA-seq analysis further associates high SPP1 expression with poor prognosis, NRF2/KEAP1 mutations and aggressive HNSCC features. In summary, our data suggest that SPP1 promotes tumor progression in NRF2- hyperactivated cisplatin-resistant HNSCC. Selective targeting of dysregulated SPP1 may offer an effective therapeutic approach for treating therapy resistant and metastatic head and neck cancers.

## Materials and Methods

### Cell Lines and cell culture

HN30 (RRID: CVCL_5525), PCI13 (RRID: CVCL_C182), PCI13 cisplatin-resistant derivatives (PCI13-wtp53, wildtype), HN30-R8, HEK293-FT (ThermoFisher, R70007), were cultured in DMEM (Gibco), containing 10% FBS, L-glutamine, sodium pyruvate, nonessential amino acids, and vitamin solution. All experiments were performed using cells from early passages routinely tested with 0.2% Myco-Zap (Lonza) to ensure mycoplasma-free culture environment (7). The HNSCC cell lines were authenticated using short tandem repeat analysis (15) within 6 months of use for the current study. The cisplatin-resistant variants HN30-R8, PCI13-wtp53 cell lines were established as previously described (7,8,14).

### Generation of SPP1 knockdown stable cell lines

The lentiviral shRNA plasmids, pGIPZ-SPP1#1 (Clone # V2LHS_303525), pGIPZ-SPP1#2 (Clone # V2LHS_303526), pGIPZ-SPP1#3 (Clone # V2LHS_303528), pGIPZ-SPP1#4 (V2LHS_111534), and pGIPZ-NT (non-targeting plasmid, Cat# RHS4348), were all obtained from Dharmacon. HN30-R8 cells stably expressing these plasmids were generated as previously described (7), and details of the experiment were described in Supplementary Materials and Methods section. HN30-R8 cells stably expressing NRF2 shRNA, wild type KEAP1 and their controls were established previously (7).

### Clonogenic survival and soft agar colony formation assays

To determine colony survival, HN30-R8 (cisplatin resistance) cells stably expressing lentiviral vector control (shCtrl) and SPP1 knockdown derivatives (shRNA SPP1 #2 and SPP1 shRNA #4) were seeded in 6-well plates and exposed to different fixed-ratios of cisplatin (CDDP; 0–20 μmol/L). Clonogenic survival and soft agar assays were performed at these concentrations as previously described by our group (16, 17).

### Western blot analysis

According to the IC50 values, minimally toxic drug doses of cisplatin (CDDP; 5-8 µmol/L) were identified from clonogenic survival assays and used for further *in vitro* experiments. Briefly, HNSCC cells and their genetically modified derivatives were treated in 10-cm dishes with CDDP (8 μmol/L) for 12, 24, and 48 hours. Whole cell lysates were prepared, and Western blot analyses were performed as described previously (16). Antibodies used for Western blotting were described in Supplementary Materials and Methods section.

### Immunoprecipitation

Whole cell lysates were prepared and collected as described previously in the western blotting method section. The immunoprecipitation was carried out using the Dynabeads™ Co- Immunoprecipitation Kit (Thermo Fisher, Cat#14321D), according to the manufacturer’s instructions. Briefly, the Dynabeads M-270 Epoxy was incubated on a roller at 37°C overnight with 5 µg antibody per mg beads for antibody coupling. The beads were washed consecutively with the washing buffer and RIPA lysis buffer. The protein lysates were incubated on a roller at 4°C for 30 minutes with the coupled beads. The beads were then washed, followed by incubation in elution buffer for 5 minutes and eluted samples were analyzed by Western blotting as described previously (16). Antibodies used for immunoprecipitation and western blotting were described in the Supplementary Materials and Methods section.

### Cell Invasion assay

Cells treated with CDDP (8 μmol/L) for 24 hours were collected, resuspended in serum-free media and seeded in the top chamber with Matrigel-coated membranes (Corning Cat # 354480). Media containing 10% FBS were added to the lower chamber. After 36 hours of incubation, the cells remaining on the upper membrane were removed with cotton wool. The cells which migrated or invaded through the membrane were stained with 20% methanol and Hemacolor (Sigma), imaged, and counted using a Leica DMLA microscope (Leica Microsystems). The experiment was carried out in duplicate wells.

### Lipid peroxidation assay

Lipid peroxidation level in ferroptotic cells was measured using BODIPY 581/59 C11 probe (Invitrogen™, D3861) in accordance with the manufacturer’s instructions. Briefly, the cells were plated in 6-cm dishes and treated with CDDP (8 μmol/l) for 48 hours as indicated. After a quick wash with PBS, the cells were incubated with 2 μmol/l of BODIPY and kept in the dark for 30 minutes at 37°C. The cells were then rinsed once with PBS to remove the staining solution, trypsinzed, and analyzed by FACS. A minimum of 100,000 cells were analyzed per dish.

### Annexin V-APC staining

HNSCC cells were seeded in 60-mm dishes, treated the next day with CDDP (8 μmol/L) and then harvested at 48 hours. Apoptosis was detected by flow cytometry using the Annexin V-APC/7-AAD staining kit obtained from BD Bioscience according to the manufacturer’s instructions.

### TCGA database analysis

Homogenized RSEM count data for the TCGA was downloaded from the TCGA TOIL database as we previously described (PMID 36002187) and upper quartile normalized to generate log2 FPKM gene expression with global median rescaling using our recently published RNA-seq pipeline (PMID 40258023). Clinical annotation and matching mutation data, including status of NRF2 and KEAP1 genes, were downloaded from the BROAD firehose web portal. Recursive partitioning to identify optimal SPP1 RNA thresholds for splitting the oral cavity squamous cell carcinoma (OCSCC) and laryngeal/hypopharyngeal squamous cell carcinoma (LHSCC) cohorts into survival groups was performed with an in-house python script that uses an iterative approach, balances groups, and maximizes significance for both Log-Rank and Gehan-Breslow-Wilcoxon approaches. Differences in SPP1 expression among groups was analyzed by a two-sided student t-test (for 2 groups), or ANOVA (for > 2 groups) with a post-hoc Tukey’s Honestly Significant Difference test. Differences in proportions of patients based on SPP1 expression group and extracapsular spread was examined with a Chi-squared test. A multiple linear regression model was fit using JMPv13 (SAS) statistical software to determine the dependence of SPP1 expression on NRF2, macrophage, and cancer associated fibroblast (CAF) single sample gene set enrichment scores (ssGSEA) and calculate standardized regression coefficients (beta values), along with associated P-values. The gene set list for the Nrf2 score was developed and validated previously by our group (PMID 36002187), while the macrophage and cancer associated gene lists were developed by vetting and refining previously published lists (PMID 31641033) using their cross-correlation coefficients of expression across more than 9,000 TCGA solid tumors as we previously described (PMID 36002187). The BROAD Gene Pattern web server tool was used to calculate individual ssGSEA values from each gene list.

### Patient samples

Following approval from the Baylor College of Medicine Institutional Review Board, archival primary and metastatic tumors from patients with a diagnosis of oropharyngeal squamous cell carcinoma, treated with curative intent at the Michael E. DeBakey Veterans Affairs Medical Center were analyzed. Patient sample characteristics are as follows: Patient #1: T3N2cM0 p16+ OPSCC treated with surgery + radiation + cisplatin, Patient #2: T2N2bM0 p16+ OPSCC treated with radiation + cetuximab, Patient #3: T2N2bM0 p16+ OPSCC treated with radiation + cisplatin, Patient #4: T2N3bM0 p16-OPSCC treated with surgery + radiation + cetuximab, and Patient #5. T4N2bM0 p16+ OPSCC treated with radiation + cisplatin (staging reflects the 7th Edition of the AJCC Staging Manual). Plasma samples for an additional 28 HNSCC patients were analyzed for SPP1 levels using standard ELISA. Patient, site and treatment characteristics for these 28 patients are summarized in Supplementary Table S8.

### SPP1 Enzyme-Linked Immunosorbent Assay (ELISA)

For detection of SPP1 levels *in vitro*, cells (1 × 10^6^) were seeded in 100-mm dishes containing complete medium and incubated for 24 hours as indicated. The cells were harvested, and the supernatants were prepared from triplicates. To measure the SPP1 *in vivo*, plasma samples from mice treated with CDDP plus anti-OPN and advanced HNSCC patients treated at Baylor College of Medicine were also obtained. The SPP1 levels were measured using the ELISA kit according to the manufacturer’s instructions (QuantikineTM, Techne Corporation R&D Systems). Briefly, cell supernatants and the plasma samples were added to the appropriate ELISA plates precoated with monoclonal antibodies directed against human OPN (SPP1). After incubation for 2 hours at 37°C, the wells in the plates were washed, and HRP-conjugated polyclonal OPN specific antibody was added. The plates were then incubated for 1 hours at 37°C, washed, and followed by addition of tetramethylbenzidine substrate. Finally, the enzymatic reactions were stopped by addition of 2 N sulfuric acid and the results were detected on a microplate reader at 450 nm.

### Reverse phase protein array (RPPA)

HNSCC cells (Ctrl and ShRNA SPP1 #4) were seeded in 10-cm dishes and harvested at 24 hours. Protein lysate was then collected from each cell line under full-serum (10% FBS) conditions and subjected to RPPA analysis as described previously (18). Detailed description of the RPPA is included in the Supplementary Materials and Methods.

### Visium Spatial Transcriptomics

Archival primary and metastatic FFPE tissue samples from the five recurrent/metastatic head neck cancer patients indicated previously in this study were processed for Visium data pre- processing and quality control to ensure high-quality data and RNA integrity. Briefly, FFPE tissue sections of 5 um thick were stained with H&E and scanned at 40X using Aperio AT2® scanner (Leica Biosystem). The H&E-stained slides underwent digital histopathology assessment using Image Scope Software X64 for pathology quality control, which included the evaluation of optimal tissue sectioning (absence of folds, tears or detachment), staining intensity and homogeneity, and scanning resolution. Only one patient (T3N2cM0 p16+ OPSCC treated with surgery + radiation + cisplatin) demonstrated optimal pathological quality and was considered for the Visium spatial transcriptomic analysis. The H&E stained slide was subjected for ST tissue processing using Visium CytAssist instrument from 10X Genomics according to the manufacturer protocol and followed by ScRNA-seq sequencing on Illumina iSeq100 for cell count validation and NovaSeq6000 at the recommended depth. To map the whole transcriptome in FFPE tissue, the reads were pre-processed using Space Ranger (v.2.1.0) from 10x Genomics and aligned to the GRCh38 reference genome. The unique molecular identifier (UMI) count matrices were analyzed using Seurat v5 in the R platform (v.4.3.0). Pathology annotations of the tissue histological morphology were performed manually by an experienced pathologist using Loupe Browser 8.0.0with the corresponding high-resolution image. Spots recognized as “excluded” which represented no tissue or damaged tissue, were filtered out during this processing step. For Visium clustering analysis, the SCTransform functions of Seurat was used to normalize and scale the matrix to identify highly variable genes (HVGs). Top 3,000 highly variable genes (HVGs) were selected for Principal Component Analysis (PCA) and downstream unsupervised spot clustering. Differentially expressed genes among clusters were identified with logFC greater than 0.25 (adjusted p < 0.05) as determined in Wilcoxon rank-sum test from Seurat. To identify spatially distinct domains, the ‘FindNeighbors’ function of Seurat was used to construct the Shared Nearest Neighbor (SNN) Graph, based on unsupervised clustering performed with ‘FindClusters’ function with a resolution set to 0.5. Subsequently, 2-D visualization of the spot clusters were performed using Uniform Manifold Approximation and Projection (UMAP) with of Seurat function RunUMAP. The top 50 PCs were used to calculate the embedding. The COMMOT (COMMunication analysis by Optimal Transport) tool was used to infer cell–cell communication analysis in spatial transcriptomics. It was installed and imported by Python following developer’s instruction. This tool which contains 1,939 validated molecular interactions and allows for detection of the interaction between different ligand and receptor species as well as spatial distances between cells (19). The analysis was performed using the standard pipeline shared by the developer. “CellChat” ligand-receptor database was used for “ct.tl.spatial communication”. The gene expression with manual masks was finally visualized using the iSTAR integrative approach which combines spatial transcriptomics with histological features (20). It is a method based on hierarchical image feature extraction that integrates ST data and high-resolution histology images to predict spatial gene expression with super-resolution. The differential gene expression in tumor cells annotated as squamous cell carcinoma by iSTAR and assigned to clusters 0, 1, 2, 7, 9, and 10 (primary tumor) or clusters 3 and 6 (metastatic lung tumor) were analyzed using the FindMarkers function in Seurat. Genes expressed in fewer than 5% of cells (p< 0.05) were excluded. An adjusted p-value < 0.01 and a log fold change > 1.5 were considered for significantly expressed genes. Gene set enrichment analyses (GESA) of these differentially expressed genes were performed using EnrichR with the 50 MSigDB HALLMARK gene sets.

### Orthotopic nude mouse model of oral cancer metastasis and therapy

All animal experimentation was approved by the Institutional Animal Care and use Committee of the University of Texas MD Anderson Cancer Center. Our orthotopic nude mouse tongue model was previously described in the literature (7). To determine *in vivo* growth kinetics and response to cisplatin treatment of HN30-R8 cells with SPP1 knockdown (SPP1 shRNA #2 and #4) and shCtrl vector, cells (5-10×10^4^) were suspended in 30 μl of medium without supplement and injected into the lateral tongues of six-week-old male athymic nude mice because oral cancer is mostly prevalent in male patients. Mice were then randomized into different groups 8 to 10 days after injection. Treatment with vehicle (PBS) or CDDP (4 mg/kg, I.V., via tail-vein/once a week for 4 weeks) was started when tumors reached a range of 3 to 6 mm^3^ in size. For the *in vivo* drug combination experiment, the HN30-R8 CDDP-resistant cells were injected into the mice oral tongues as described above, and treated with either CDDP alone, the SPP1 inhibitor (osteopontin monoclonal antibody, BioXCell #Cat #BE0382, 10 mg/kg, 3 times a week, given intraperitoneally, I.P.) alone, and their combination or PBS vehicle (given I.P.) for 4 weeks. A total of 10 to 12 mice in each group were used, and 90% of the mice had tumor growth under each condition. Mice were monitored twice a week, and their weight and tumor volume were recorded. Tongue tumors were measured in millimeters with microcalipers by investigator blinded to the treatment groups, and tumor volume was calculated as A × B2 × (π/6), where A is the longest dimension of the tumor and B is the dimension of the tumor perpendicular to A. Mice were euthanized when they lost more than 20% of their preinjection body weight. During necropsy, tongues, cervical lymph nodes and lungs were harvested, formalin-fixed, and subjected to histologic evaluation using hematoxylin and eosin (H&E) staining to identify primary tumors and count the number and size of spontaneous metastases. A separate group of mice were injected with HN30-R8 cells expressing shCtrl vector and SPP1 shRNA#2 via tail vein injection to confirm incidence of spontaneous metastasis. Lungs were harvested 60 days after tail injection of the cells, processed and evaluated with H&E staining as described above.

### Statistical analyses

The student’s t and two-way ANOVA tests were utilized to analyze *in vitro* and *in vivo* data. Mice survival following tumor formation or drug treatment was analyzed by the Kaplan–Meier method and compared with log-rank test. Student t-test statistic value of P<0.05 was used to compare ELISA results in mice and human plasma samples. All data were expressed as mean ± standard error, and cutoff p-values of 0.05 or less were considered to indicate statistical significance. *In vitro* experiments were carried out in triplicate or duplicate as appropriate (for each condition) and were repeated to ensure reproducibility.

### Data availability statement

The data generated in this study are available within the article and its Supplementary Data files. Data used in this study that is not included in the paper or supplementary files can be made available upon request from the corresponding author.

## Results

### SPP1 targeted suppression enhances cisplatin sensitivity *in vitro* in cisplatin resistant HNSCC through induction of ferroptosis

To investigate whether Nrf2-activated SPP1 contributes to cancer cell growth and cisplatin resistance in HNSCC, we initially evaluated the SPP1 expression levels in cisplatin sensitive (HN30) and resistant (HN30R-8) cell lines following treatment with cisplatin (CDDP) as indicated. Western blot analysis showed that acquired cisplatin resistance was associated with increased SPP1 protein levels in HN30-R8 cells compared to HN30 cells **(Fig. 1A).** Restoration of wt*KEAP1* or loss of NRF2 is accompanied by reduction in SPP1 protein levels in HN30-R8 cells, suggesting that SPP1 is one of the primary downstream targets modulated by dysregulated KEAP1/NRF2 signaling axis during acquired cisplatin resistance in HNSCC **(Fig. 1B).** We next utilized SPP1 shRNA and control lentiviral vectors to generate stable SPP1-knockdown HN30- R8 and PCI13-wtp53R cell lines **(Fig. 1C, Supplementary Figure S1A and S1B) r**espectively. Western blot analysis demonstrated successful knockdown of SPP1 in these cell lines and HN30-R8 cells with SPP1 shRNA #2 and #4 were considered for further analysis **(Fig. 1C).** Because SPP1 is a secretory protein, we confirmed its level by ELISA in conditioned media prepared from these cells (Fig. 1D). Suppression of SPP1 enhanced cisplatin sensitivity and reduced the CDDP IC_50_ from 8.48 μmol/L in cells with lentiviral control (shCtrl) vector to 5.23 and 3.34 μmol/L in cells with lentiviral shRNA SPP1 #2 and #4 vectors respectively **(Fig. 1E and F).**

**Figure 1.**
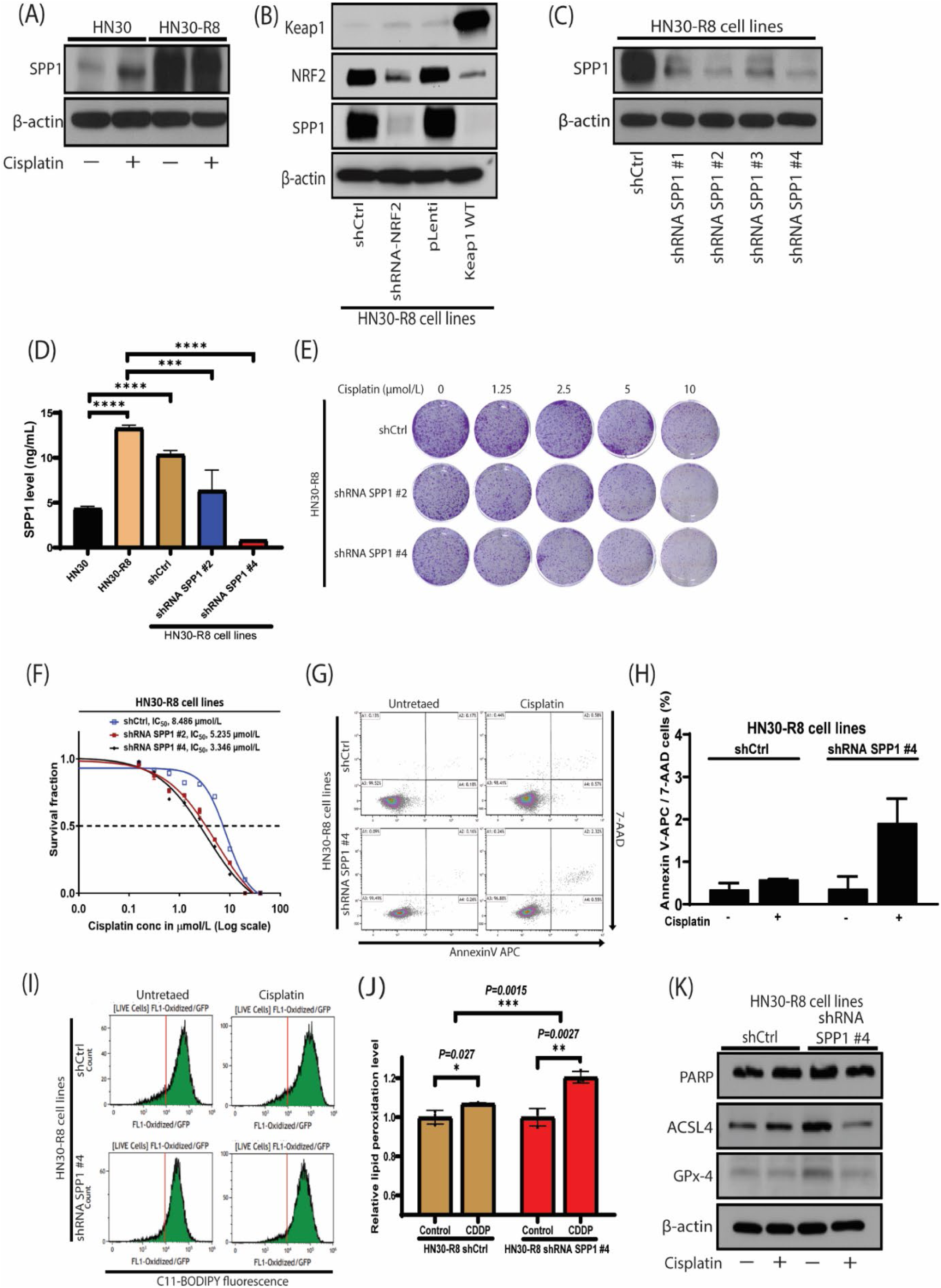
Downregulation of SPP1 enhances cisplatin sensitivity *in vitro* in cisplatin resistant HNSCC through induction of ferroptosis. **A,** Western blot demonstrates high expression of SPP1 during acquired CDDP resistance in HN30-R8 HNSCC cells compared to HN30 CDDP sensitive parental cells. **B,** Restoration of KEAP1 or knockdown of NRF2 modulates SPP1 expression in CDDP resistant HNSCC cells. **C,** Western blot verifies downregulation efficiency of SPP1 in stable CDDP resistant cells (HN30-R8). **D,** Bar graph depicting levels of SPP1 determined by ELISA in CDDP sensitive (HN30), resistant (HN30- R8) and their SPP1 knockdown derivatives. **E** and **F,** Representative images of clonogenic survival and curves demonstrating increased sensitivity to CDDP in HN30-R8 cells stably expressing shRNA SPP1 after treatment with the indicated doses of cisplatin. **G** and **H,** Annexin V-APC/7-AAD–positive staining (percentage of dead cells) confirming induction of low levels of apoptosis in SPP1 knockdown HN30-R8 cells treated with CDDP (8 µmol/L) for 48 hr. **I,** Fluorescence level of intracellular oxidized C11-BODIPY (581/591) measured by Flow cytometry. **J,** Bar graph showing increased lipid peroxidation levels calculated from the fluorescent integrated density of the oxidized BODIPY indicates ferroptosis in HN30-R8 cells expressing SPP1 shRNA following treatment with CDDP (8 mol/L) for 48 hr compared to untreated ShCtrl control cells. **K,** Western blot demonstrates decreased protein level of anti- ferroptotic markers, ACSL4 and GPX4 and absence of apoptotic marker, PARP cleavage. Data shown are representative of three independent experiments. Bar graphs are mean ± SEM, unpaired student t-test and two-way ANOVA respectively. **P < 0.001, ***P< 0.0001, *P=0.027, **P=0.0027, ***P=0.0015.

Sensitivity to cisplatin was also observed in the PCI13-wtp53R cells expressing shRNA SPP1 #2 and #4 compared to the parental HN30 and shCtrl cells **(Supplementary Figure S1C and S1D).** This result is consistent with published data demonstrated that SPP1 increased cisplatin resistance in lung cancer cells (21). To determine how suppression of SPP1 sensitized cells to CDDP, mechanisms of cell death were assessed. SPP1 depletion has minimal effect on apoptosis **(Fig. 1G and H**). We next evaluated if suppression of SPP1 increased lipid peroxidation and induced ferroptosis in CDDP resistant HNSCC cells. Compared to control cells, HN30-R8 cells expressing shRNA SPP1 had a shift in excitation and emission upon oxidation (**Fig. 1I**) and significantly higher levels of lipid peroxidation upon CDDP exposure **(Fig. 1J)**, indicating cell death through ferroptosis. Furthermore, the lack of cleaved PARP1, a marker of apoptosis, and the decreased protein levels of the inhibitory ferroptosis markers, GPX4 and ACSL4 determined by western blot analysis confirmed induction of ferroptosis in the cells tested **(Fig. 1K).**

### Knock-down of SPP1 reduces *in vitro* colony growth and invasion in HNSCC cisplatin- resistant cells

Since SPP1 has been shown to promote tumor cell migration and invasion (9), colony formation and transwell migration assays were performed as previously described, to examine whether SPP1 downregulation exerted similar effect on CDDP-resistant HNSCC cancer cells. Compared to cells with lentiviral vector control (HN30-R8 shCtrl), the colony growth was significantly reduced in HN30-R8 shRNA SPP1 and this inhibitory effect is further enhanced upon treatment with CDDP (8 μmol/L) **(Fig. 2A and B).** Suppression of SPP1 inhibited cell invasion in both untreated and treated HN30-R8 cells (P<0.0001) **(Fig.2 C and D).** Similar results were obtained with anti-osteopontin antibody treatment of HN30-R8 cells **(Fig. 2E and F).**

**Figure 2.**
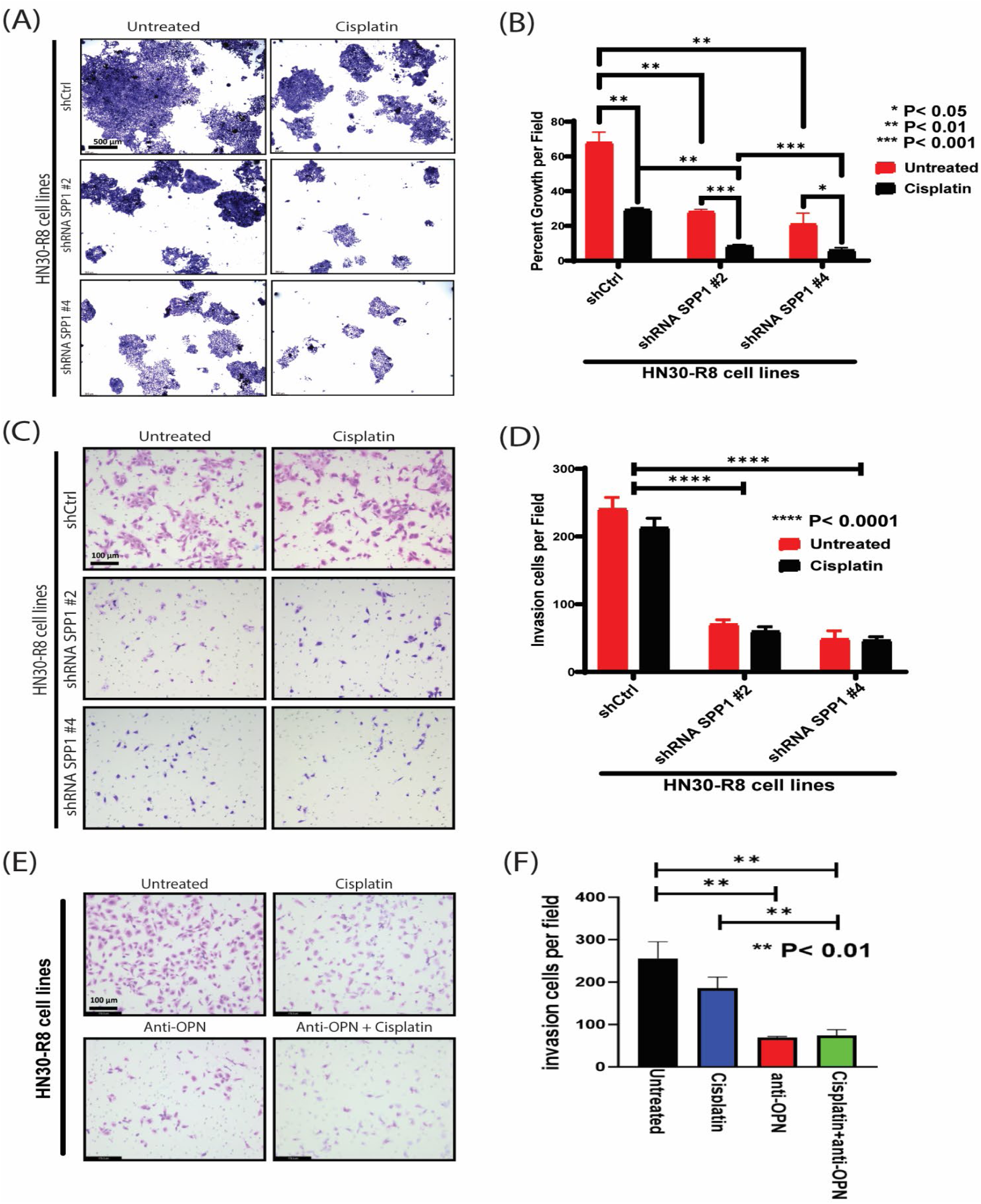
Inhibition of SPP1 reduces *in vitro* colony growth and inhibits cell invasion in HNSCC cisplatin-resistant cells. The HN30-R8 and their shRNA SPP1 derivatives were cultured in presence and absence of CDDP (8 mol/L) for 48 hr and subjected to colony growth on soft agar and Matrigel invasion assays as described in Methods**. A and B,** Knock-down of SPP1 alone or with CDDP treatment decreases the colony growth on soft agar. **C and D,** The invasion ability of HN30-R8 cells expressing shRNA SPP1 is significantly decreased compared to the shCtrl control cells. **E and F,** Cell invasion is inhibited following neutralization of the SPP1 with the anti-SPP1(osteopontin) antibody and in the presence of CDDP in HN30-R8 CDDP resistant cells. Data shown are representative of three independent experiments. Error bars are mean ± SEM, unpaired student t-test and two-way ANOVA respectively. *P<0.05, *P<0.01, ***P<0.001, ****P < 0.0001. The scale bar is 100 µm. Images were taken using Leica DMLA microscope.

### SPP1 targeted suppression potentiates sensitivity to cisplatin and inhibits tumor growth and metastasis *in vivo* in cisplatin-resistant oral cancer model

To evaluate the impact of altered SPP1 on CDDP sensitivity *in vivo*, we injected the HN30-R8 stably expressing shRNA SPP1 #4 and shCtrl vectors into the tongues of athymic nu/nu mice followed by CDDP treatment as indicated. As expected, no significant response was observed upon CDDP treated bearing HN30-R8 shRNA Ctrl transfected cells **(Fig. 3A).** Consistent with published data (22), tumor growth in untreated mice orally injected with cells expressing shRNA SPP1 #4 vector was slightly decreased compared to the untreated shRNA Ctrl control group **(Fig. 3A).** However, significant tumor growth inhibition was observed following CDDP treatment in mice injected with shRNA SPP1 #4 cells, compared with all other treated or untreated groups (P < 0.0001; **Fig. 3A).** CDDP treatment was associated with improved animal survival in mice bearing shRNA SPP1 #4 cells as compared to the untreated group and animals in the CDDP treated and untreated HN30-R8 shRNA controls (P = 0.0017; **(Fig. 3B).** Body weight was not significantly different between the groups with or without CDDP treatment **(Fig. 3C).** We assessed the effect of SPP1 down-regulation on the development of lung metastases of CDDP resistant HNSCC cells *in vivo*. Mice were injected orally with the shCtrl and ShRNA SPP1 #4 cells followed by treatment with CDDP as indicated. A total of 12 mice (6 untreated and 6 CDDP treated) were used in each group. Histological H&E staining was utilized to evaluate for the presence of microscopic metastases in the lymph node and lung sections obtained from these mice. We defined N0 as non-metastatic, N1 as simple metastasis (metastasis to one lymph node, and N2 as multiple metastases (metastases to more than one lymph) in cervical lymph node **(Fig 3D-F and Table 3G.** Mice harboring HN30-R8 shCtrl tumors had higher rates of cervical lymph node metastases (9 mice with N2 score), and 5 of the 9 mice with staged as N2 in this group had multiple lung metastases **(Fig. 3D-F and Supplementary Figure S2A).** Mice harboring tumors with SPP1 knockdown had fewer number N2 (2/12 mice) metastases **(Fig. 3D- F and Supplementary Figure S2A).** None of the mice with SPP1 knockdown were found to have lung metastasis **(Fig. 3F and Table 3G).** Additionally, the accelerated rate of distant metastasis was further confirmed in the tail vein metastatic model using tumor cells expressing the shCtrl vector (8 mice) and other shRNA SPP1 construct #2 (8 mice) **(Fig. 3H).** While all mice in the control group (shCtrl) had lung metastases, only two mice in the shRNA SPP1 #2 group were found to have a very low number metastatic nodules in their lungs (**Fig. 3I-J, Table 3K and Supplementary Figure S2B).** A significant improvement in mouse survival (p = 0.0085) was observed in the SPP1 knock down group as compared to the HN30-R8 shRNA. Taken together, these data suggest that dysregulated SPP1 plays a critical role in tumor progression and metastasis in NRF2 activated cisplatin resistant HNSCC.

**Figure 3.**
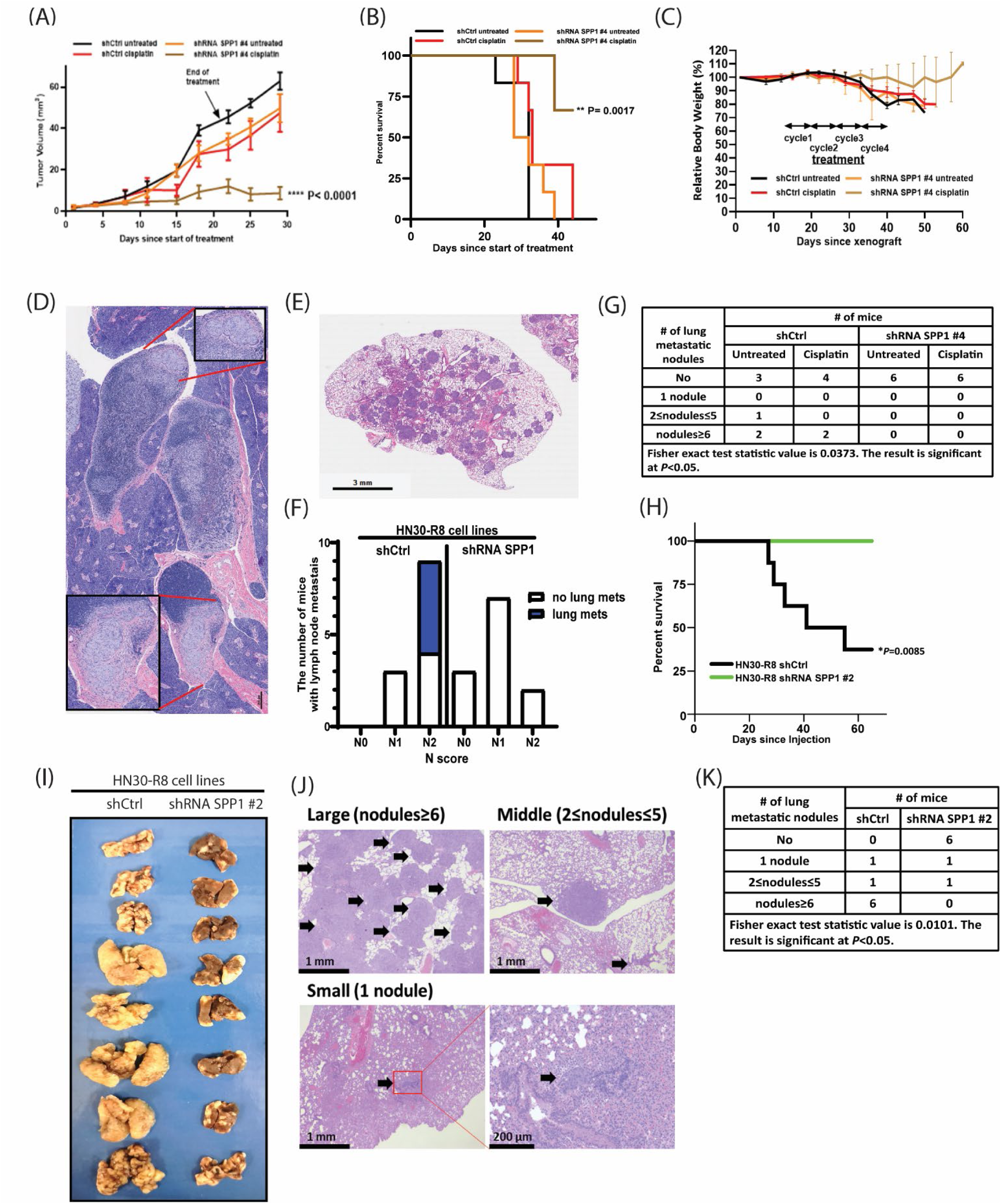
SPP1 targeted suppression enhances cisplatin sensitivity and reduces tumor growth and metastasis in vivo in cisplatin resistant oral cancer model. HN30-R8 cells stably expressing shRNA SPP1 #4 and shCtrl constructs were orthotopically injected into the tongues of male athymic nu/nu mice and treated intravenously via tail injection with 4 mg/kg of CDDP for 4 weeks as indicated. Each treatment group contains 9-12 mice. Tumor growth was measured and reported as tumor volume means ± SEM. A, Tumor growth curves calculated after 4 weeks of injection and treatment. Statistical analyses were performed by a two-way ANOVA test. ****P < 0.0001 CDDP treated HN30-R8 shRNA SPP1 #4 tumor bearing mice versus all other treatment groups. B and C, Kaplan–Meier survival curve and percent of body weight loss of mice from A. The death of animals occurred when tumor compromised the animal welfare. D and E, Incidence of lymph node and lung tumor metastases in mice assessed microscopically by hematoxylin and eosin staining. F, Bar graph depicting the impact of SPP1 knockdown on the number of lymph node (N score) and lung tumor metastases present concurrently in mice. G, Table summarizes the number of mice with lung metastatic nodules, where No (indicates no nodule present). H, Kaplan–Meier survival curve of mice injected with HN30-R8 (shCtrl) and HN30-R8 (shRNA SPP1 #2 construct) cells through the tail vein. I, Macroscopic nodules can be seen in the representative images from the lungs of mice injected with the tumor cells via tail vein. J and K, respective microscopic images and Table depicting the number of the lung metastatic nodules, and their sizes appearing in mice following tail vein injection with the tumor cells. All in vivo data were expressed as ± SEM and Log-rank, *P< 0.05 was considered significant for mice survival. The Fisher exact test statistic value, *P<0.05 was used to detect significant difference in lymph node and lung metastasis in mice.

### Pharmacological inhibition of SPP1 enhances cisplatin sensitivity *in vivo* in CDDP resistant oral cancer model

As SPP1 suppression led to significant tumor growth reduction and improved sensitivity to cisplatin *in vivo,* we sought to pharmacologically target SPP1 and evaluate its impact on tumor growth and metastasis using a commercially available anti-osteopontin (SPP1) antibody. HN30- R8 tumors were randomly assigned to one of 4 treatment groups: 1) control, 2) CDDP, 3) anti-OPN antibody and 4) anti-OPN antibody in combination as indicated in the protocol outlined in **Figure 4A**. Antibody dose (10 mg/kg) and frequency of administration was chosen based on the *in vitro* data **(Fig. 2E)**. No significant anti-tumor efficacy was observed with the anti-OPN treatment alone at this selected dose compared to either untreated or CDDP treated groups. However, the combination of CDDP and anti-OPN antibody was associated with delayed tumor growth and improved animal survival **(Fig. 4B and C**). No significant body weight loss was seen among the treatment groups, indicating that the combination was well tolerated **(Fig. 4D).** Plasma levels of OPN (SPP1) were significantly reduced after treatment with anti-OPN antibody given alone or in combination with CDDP compared to untreated mice or those treated with CDDP alone **(Fig. 4E).** OPN (SPP1) levels were decreased in the anti-OPN and combination treatment groups to a level comparable to that obtained from mice implanted with the CDDP sensitive parental HN30 cell line **(Fig. 4E).** These data confirmed that the anti-OPN dose used was able to deplete SPP1 and prevent binding to its cognate receptors *in vivo*. Additionally, the effect of anti-OPN antibody and its combination with CDDP on tumor progression and metastasis was microscopically evaluated in the mice lymph nodes and lungs sections stained with H&E as indicated. Consistent with published report (23), only the combination of anti-OPN antibody and CDDP was able to reduce the number of mice with cervical lymph nodes (1 out of 6 mice; 17%) and lung metastasis (3 out of 6 mice; 50%) compared to other treatment groups **(Table 4F and Supplementary Figure S3A and B),** suggesting that OPN (SPP1) plays an important role in tumor growth and regional and distant metastasis.

**Figure 4.**
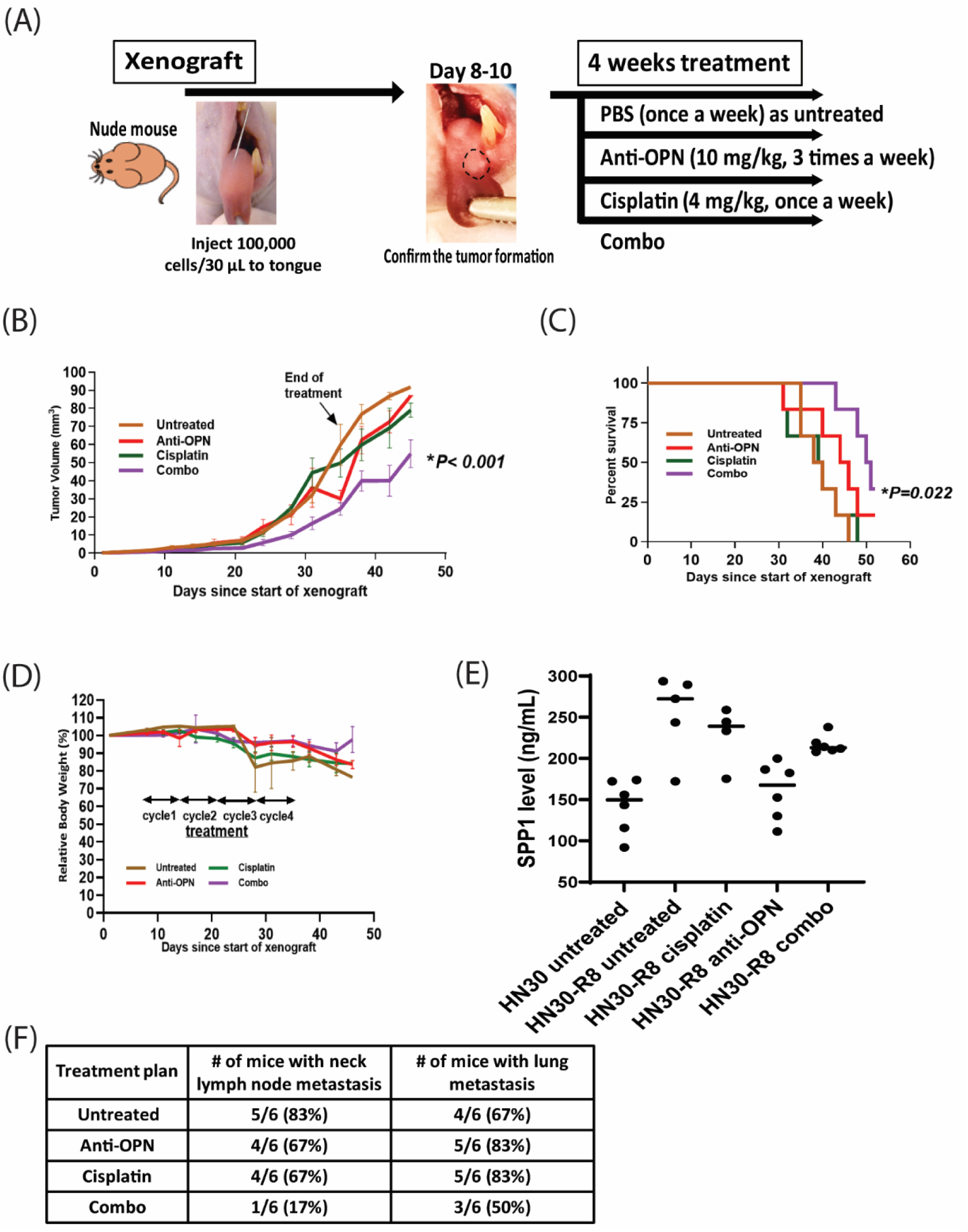
Pharmacological targeting of SPP1 improves cisplatin sensitivity and reduces tumor growth *in vivo* in CDDP resistant oral cancer model. **A)** Schematic procedure of the *in vivo* mouse xenograft experiment and treatment schedule. Mice were injected with CDDP resistant HN30-R8 cells and after tumor formed, mice were treated with CDDP, anti-OPN, and in combination as indicated. **B,** Tumor growth curves calculated after 4 weeks of injection and treatment with drugs. **C,** Kaplan–Meier survival curve demonstrating improved survival of mice following combination of anti-OPN and CDDP. **D,** Percent body weight loss among the treatment groups. **E,** SPP1 levels determined by ELISA-based assay in plasma prepared from mice blood samples collected following treatment with drugs. The plasma samples were diluted 100 times in the ELISA dilution buffer. **F,** Table showing microscopic evaluation of incidence and percentage of tumor metastatic nodules in mice lymph nodes and lungs sections stained with H&E as indicated. All *in vivo* data were expressed as ± SEM and two-way ANOVA and Log-rank tests, *P< 0.05 were considered significant for tumor growth reduction and mice survival, respectively.

### SPP1 gene expression is a poor prognostic factor that correlates with *NRF2/KEAP1* mutational status and aggressive clinical features in HNSCC

We analyzed RNAseq data from the TCGA HNSCC cohort to investigate how SPP1 gene expression correlates with clinical features and prognosis. HNSCC tumors were initially stratified by subsite—including oral cavity SCC (OSCC), laryngeal/hypopharyngeal SCC (LHSCC), and oropharyngeal SCC (OPSCC) as well as by HPV status, to compare SPP1 expression with adjacent normal samples. Compared to normal samples, levels of SPP1 expression were significantly increased in tumors from nearly all the subsites, regardless of HPV status, with the exception of HPV-positive LHSCC **(Fig. 5A).** Next, we examined whether mutations in either *NFE2L2* or *KEAP1* were associated with increased SPP1 expression because SPP1 is a known downstream target of NRF2 activation. Collectively, tumors with a mutation in either one of these genes had significantly elevated average SPP1 mRNA (P<0.001, **Fig. 5B)** in the OCSCC cohort, and a similar trend was found among LHSCC tumors that nearly reached significance. OCSCC and LHSCC patients with high SPP1 expression in their tumors, determined through recursive partition analysis, had significantly reduced overall median survival times **(Fig. 5C and D)**, which were also associated with reduced median disease-free survival times **(Fig. 5E and F)**. Consistent with these findings, OCSCC tumors from patients with a higher N stage (>N1) and T stage (T3 &T4) had significantly increased average expression of SPP1 **(Fig. 5G and H),** and reciprocally patients with tumors categorized as high SPP1 expression were associated with an increased frequency of gross (e.g., Macro) extracapsular spread **(Fig. 5I).** Likewise, tumors associated with the presence of gross extracapsular spread had a tendency towards increased SPP1 expression **(Fig. 5J).** The average SPP1 expression of poorly and moderately differentiated tumors was increased relative to well differentiated tumors, with the latter more than two-fold higher (P=0.009, **Fig. 5K).** Because SPP1 can be expressed by macrophages, cancer-associated fibroblasts (CAF), tumor cells, and possibly other cells present in the tumor microenvironment we examined the putative relationships between SPP1 expression, the presence of various cell types, and survival. To untangle the possible connections between SPP1 expression and cell types in the tumor microenvironment we first performed single sample gene set enrichment analysis (ssGSEA) on TCGA OCSCC samples using published gene lists specific for enrichment of 19 different cell types that included various leukocyte subsets, endothelial cells, CAF, and cross correlated the scores with SPP1 mRNA expression and also NRF2 activation scores **(Supplementary Table S1).** Hierarchical clustering of the cross-correlation coefficients enabled visualization of feature modules that behaved similarly **(Fig. 5L),** where we observed high co-correlation among a subset of leukocytes including eosinophils, neutrophils, monocytes, mast cells, and macrophages―all moderately correlated with SPP1 levels. Among these we chose macrophages to represent this leukocyte group in building a multiple linear regression model of SPP1 expression because of prior literature documenting SPP1 expression in macrophages. We also examined NRF2 scores and CAF scores as additional predictors because they appeared to independently correlate with SPP1 from the heatmap. The regression model **(Fig. 5M)** revealed that NRF2 was the strongest predictor of SPP1 (β = 1.11, p < 0.00001), followed by macrophages (β = 0.82, p < 0.00001), and CAF (β = 0.60, p < 0.00001). Finally, we performed univariate and multivariate cox regression considering SPP1, macrophage score, NRF2 score, and CAF score as predictors of overall survival using the TCGA OCSCC cohort **(Supplementary Table S2).** SPP1 was the only variable to correlate with overall survival in either the univariate (β = 0.237, HR = 1.27, P = 0.0078) or multivariate (β = 0.277, HR = 1.32, P = 0.0078), indicating that high SPP1 expression was an independent significant predictor of poor overall survival.

**Figure 5.**
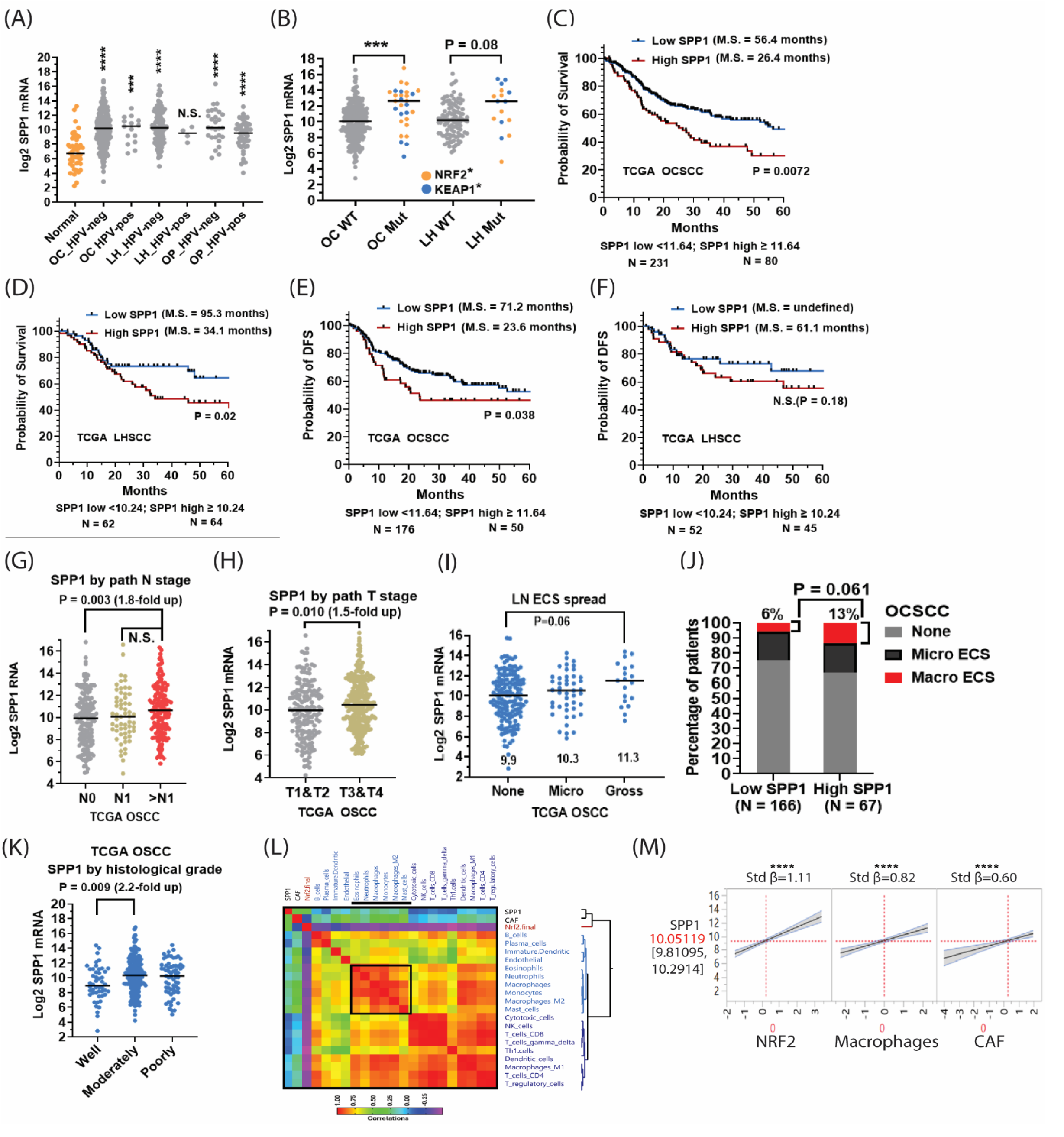
SPP1 correlates with poor prognosis, *NRF2/KEAP1* mutational status, and aggressive clinical features in HNSCC. **A,** Levels of SPP1 mRNA are significantly higher in oral cavity (OC) squamous cell carcinoma (OCSCC) and laryngeal / hypopharyngeal (LH) squamous cell carcinoma tumors compared to normal samples from the TCGA cohort, regardless of HPV association. **B,** SPP1 expression is elevated in tumors harboring mutations in either NRF2 (orange symbols) or KEAP1 (blue symbols) genes. **C,** OCSCC tumors with elevated SPP1 mRNA expression (cohort cutoff determined by recursive portioning) have significantly lower median overall survival time. **D,** LHSCC tumors with elevated SPP1 mRNA expression (cohort cutoff determined by recursive portioning) have significantly lower median overall survival time. **E,** OCSCC tumors with elevated SPP1 mRNA expression have significantly lower median disease-free survival (DFS) times. **F,** LHSCC tumors with elevated SPP1 mRNA expression have significantly lower median DFS times. **G,** OSCC patients with pathological lymph node stage >N1 have significantly higher SPP1 mRNA expression compared to node negative (N0) patients. **H,** OCSCC patients with more advanced T stages (T3 & T4) have significantly elevated SPP1 mRNA expression. **I,** Primary tumors with higher levels of SPP1 mRNA had a greater incidence of gross extracapsular lymph node extension. **J,** Primary OCSCC tumors associated with microscopic (micro) or gross extracapsular lymph node spread had a trend towards elevated SPP1 mRNA expression. **K,** Elevated expression of SPP1 mRNA was associated with less differentiated OCSCC tumors. *P ≤ 0.05, **P ≤ 0.01, ***P ≤ 0.001, ****P ≤ 0.001. **L,** Hierarchical clustering of the cross-correlation coefficients featuring modules highly co-correlated among a subset of leukocytes including eosinophils, neutrophils, monocytes, mast cells, and macrophages which all moderately correlated with SPP1 expression levels. **M,** Regression model revealing that NRF2 is the strongest predictor of SPP1 (****p < 0.00001), followed by macrophages. (****p < 0.00001), and CAF (****p < 0.00001).

### Reverse phase protein array (RPPA) analysis identifies protein analytes and metastatic signaling pathways downstream of SPP1 expression

Binding of SPP1 to its receptors triggers a cascade of intracellular signaling pathways that regulate cell proliferation and tumor progression. Therefore, to investigate the signaling pathways by which Spp1 promotes tumor progression in CDDP resistant HNSCC tumors, we employed reverse phase protein array (RPPA) analysis to identify changes in total and phospho- protein levels for 77 analytes in HN30R8 cells after disrupting SPP1 expression with shRNA. Significant changes were found for 33/77 analytes after SPP1 shRNA knockdown **(Supplementary Table S3 and Fig. 6A),** and all analytes decreased in level after SPP1 knockdown compared to controls shRNA **(Fig. 6A and B).** Gene ontology pathway enrichment using the list of 33 altered analytes identified a number of enriched tumor promoting processes including immune response, stress response, differentiation, cell migration, MAPK signaling cascade, and AKT/mTOR signaling **(Supplementary Table S3 and Fig. 6B).** Notably, after silencing SPP1 in HN30R8 we observed decreases in protein expression of pAKT (T308), pFAK1 (Tyr397), pGSK3α-β-(pS21-S9), p-p90RSK (Thr573), and p-PAK1 (Thr423/PAK2 (Thr402) respectively. Downregulation of these proteins was further confirmed by western blotting **(Fig. 6C).** Increased levels of these phosphorylated proteins are associated with tumor cell survival and metastasis in several cancer types, making them potential protein targets in HNSCC. These data suggest that SPP1 inhibition causes significant decrease in relevant oncogenic and metastatic signaling pathways and functions during cisplatin resistant in HNSCC.

**Figure 6.**
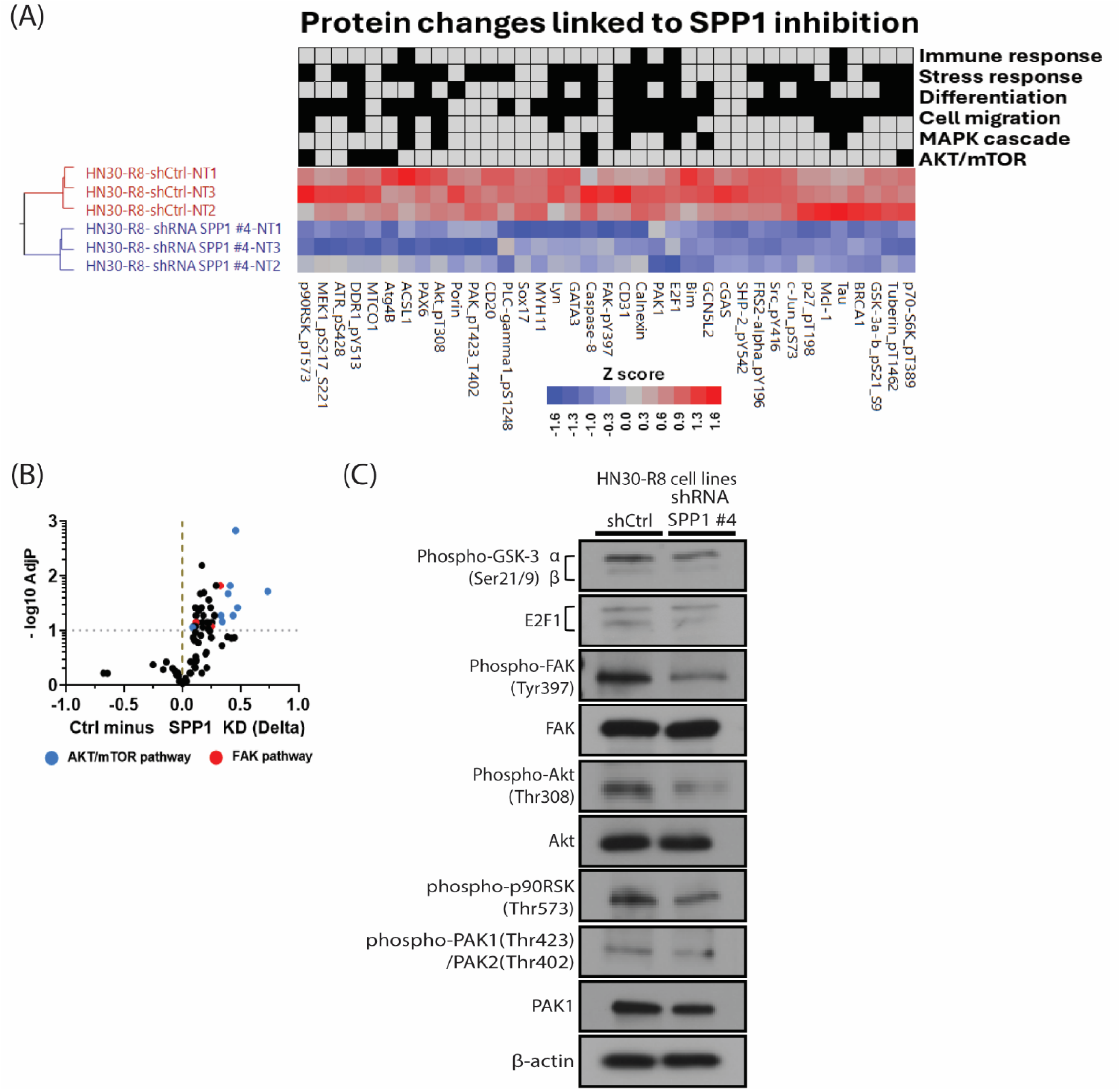
Reverse phase protein array (RPPA) analysis identifies protein analytes and metastatic signaling pathways downstream of SPP1 expression. **A,** Volcano plot showing negative log10 adjusted P values (y-axis) verses differences in RPPA values (x-axis) measured for analytes derived from HN30R8 after infection with either control shRNA (shCtrl) or shRNA SPP1 knockdown (SPP1 KD). Positive delta values indicate analytes that decrease after SPP1 KD compared to control. Analytes in the AKT / mTOR pathway are represented with blue symbols, while analytes from the FAK pathway are represented with red symbols. A doted horizontal line at Y =1 corresponds to a minimum false discovery rate of 0.1. **B,** Heatmap of select analytes measured by RPPA after one-way unsupervised hierarchical clustering, showing that samples clustered by the status of SPP1 KD. Gene Ontology (GO) enrichment was performed on all analytes found significantly different after SPP1 KD and analytes from select GO pathways found to be significantly enriched were included in the cluster analysis and are annotated by GO process with black squares above the heatmap. **C,** Western blot analyses confirming differential expression of most of these proteins, including AKT/ mTOR and FAK among the control and SPP1 KD cell lines.

### Visium spatial transcriptomic analysis in metastatic HNSCC tumor reveals a potential mechanistic interplay between tumor-derived SPP1, integrins and CD44 receptors

The continuous interplay between tumor cells and the tumor microenvironment (TME) strongly affects tumor development, disease progression, metastasis, and responses to therapeutic interventions. The spatial composition and microenvironmental ecological niche within tumors have been reported to be associated with TME remodeling and tumor metastasis (24). In this study we have shown that elevated level of SPP1 is associated with the development of cisplatin resistance, tumor progression and metastasis *in vivo* in NRF2-hyperactivated HNSCC. This is significant because recent studies have highlighted the pivotal role of SPP1 in the progression of malignant tumors through modulation of TME (25,26). Therefore, we sought to investigate whether the metastatic potential of cisplatin-resistant HNSCC is linked to a spatial interaction between NRF2-activated SPP1 signaling and the TME. We performed Visium spatial transcriptome (ST) on archival primary and metastatic FFPE tissue sample from one recurrent/metastatic head neck cancer patient. H&E staining and UMAP revealed distinct, color-coded cell types and pathologically annotated clusters (spots) from the ST dataset of primary and metastatic lung tumors. **(Fig. 7A).** As expected, the squamous cell carcinoma represented the majority of the cell type in these tumors **(Fig. 7A).** The top 3,000 highly variable genes (HVGs) were selected for Principal Component Analysis (PCA) and downstream unsupervised spot clustering. The iSTAR analysis in tumor samples identified a KEAP1/NRF2 gene signature in 400 out of over 1000 genes **(Supplementary Table S4).** Although, KEAP1 and NRF2 expression was similar in primary and metastatic tumors, the SPP1 expression was significantly higher in lung metastatic tumors compared to the primary tumor **(Fig. 7B).** This increased SPP1 expression is consistent with its known functional role in promoting tumor metastasis. SPP1 acts through binding to various cell surface integrins and CD44 receptors and activates downstream signaling pathways to promote tumor growth, invasion, and chemoresistance (27). However, its interaction with these receptors during cisplatin resistance or tumor progression in HNSCC is not fully elucidated. COMMOT is a computational method which incorporates the spatial information and ligand-receptor interactions in spatial transcriptomic data. Here, we combined the iSTAR with the COMMOT to identify spatial cell-cell communication patterns in the HNSCC patient tumors from the gene expression, focusing on SPP1 and its cognate receptors. High expression levels of major SPP1 receptors including CD44 and several integrin subunits (α4, αv, β1, β3, and β6) were observed in the cell types in the patient primary and metastatic lung tumors **(Fig. 7C),** revealing a strong interaction between SPP1 and several integrins (α4β1, αvβ1, αvβ3, and αvβ6) in both primary and metastatic tumors **(Fig. 7D**). In HNSCC lung metastases, SPP1 appears to bind specifically to CD44, integrin αvβ1 and integrin αvβ6 (**Fig. 7D**). Immunoprecipitation analysis on CDDP-resistant HN30-R8 cell lines *in vitro* partly confirmed this interaction, showing strong co-immunoprecipitation of SPP1 with integrins α4β1, αvβ1, and αvβ3, but not with CD44, in CDDP-resistant HN30-R8 cells **(Fig. 7E).** This validation clearly supports the interaction of SPP1 with its predicted integrin receptors, as identified by spatial transcriptomic and COMMOT analysis.

**Figure 7.**
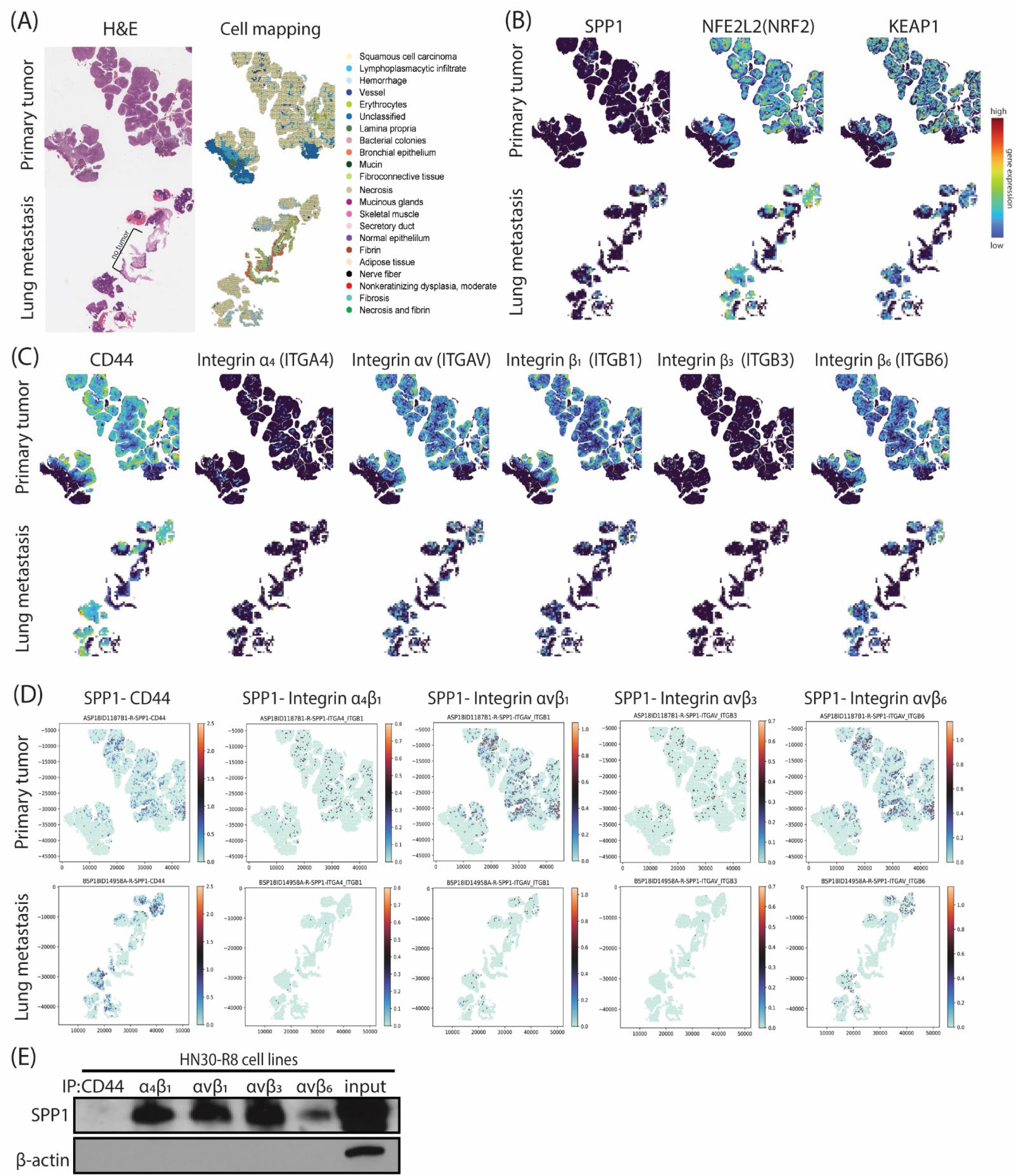
Visium spatial transcriptomic analysis in metastatic HNSCC reveals a potential mechanistic interplay between tumor-derived SPP1, integrins and CD44 receptors. A, Hematoxylin and eosin staining and UMAP plots showing the histological annotation of the H&E -stained Visium slides and cell types in patient primary and metastatic lung tumors. **B,** Near super-pixel resolution analysis using iSTAR and visualization of differential expression of SPP1, NFE2L2 (NRF2), KEAP1 and KEAP1 in primary and metastatic lung tumors. **C,** Near super-pixel resolution analysis using iSTAR and visualization of the differential expression of SPP1 receptors in primary and metastatic lung tumors. **D,** COMMOT clustering analysis and similar communication intensity signals demonstrating possible interaction between SPP1 and its receptors, CD44 and integrins in primary and metastatic lung tumors in HNSCC patient. **E,** Western blot analysis showing positive interaction of SPP1 with CD44 and several integrin receptors *in vitro* in CDDP-resistant HN30-R8 cell lines.

### Spatial annotation and gene set enrichment analyses identify high expression of SPP1 and distinct cancer HALLMARK pathways between primary and metastatic lung tumors in HNSCC patient

To analyze gene expression differences between paired primary and metastatic lung tumors in HNSCC patient, differentially expressed genes were identified within 14 cell clusters (cluster 0-13) using Seurat **(Fig.8A).** The pathology annotation of the cell types and clusters in the primary and metastatic lung tumors was analyzed by Seurat **(Supplementary Table S5).** These genes were then annotated to understand their function and roles within each cluster. This approach allows for a detailed comparison of molecular changes occurring during tumor metastasis. The top 6 significantly expressed genes in each cluster were presented as bubble and annotation clustering plots **(Supplementary Figure S4).** Our data further showed that clusters 0, 1, 2, 7, 9, and 10 were more prevalent in the primary tumor, while clusters 3 and 6 were the predominant features of the metastatic lung tumor **(Supplementary Table S5)**. Notably, the SPP1 expression was higher in the metastatic lung tumor **(Fig. 8B)**. Only 525 genes with logFC>1.5 and FDR <0.01 were considered for further analysis. Therefore, 186 differentially expressed genes were significantly upregulated in the primary tumor and 339 genes in the metastatic lung tumor as shown in the volcano plot **(Fig. 8C and supplementary Table S6)**. On the basis of these findings, we next performed functional HALLMARK and enrichment analyses of these target genes using the AUCell package to identify different gene signatures active in cells from primary and metastatic lung tumors **(Fig. 8D)**. Primary tumor clusters showed significant enrichment of genes related to epithelial-mesenchymal transition (EMT) signatures, while metastatic lung tumor clusters exhibited enrichment of interferon response signatures **(Fig. 8E and supplementary Table S7A and S7B)**. Collectively, our findings suggest that SPP1 interacts with its known cognate receptors, including integrins and CD44 in HNSCC tumors. This interaction is hypothesized to enhance oncogenic cellular responses, eventually contributing to therapy resistance, tumor progression and metastasis **(Fig. 8F).**

**Figure 8.**
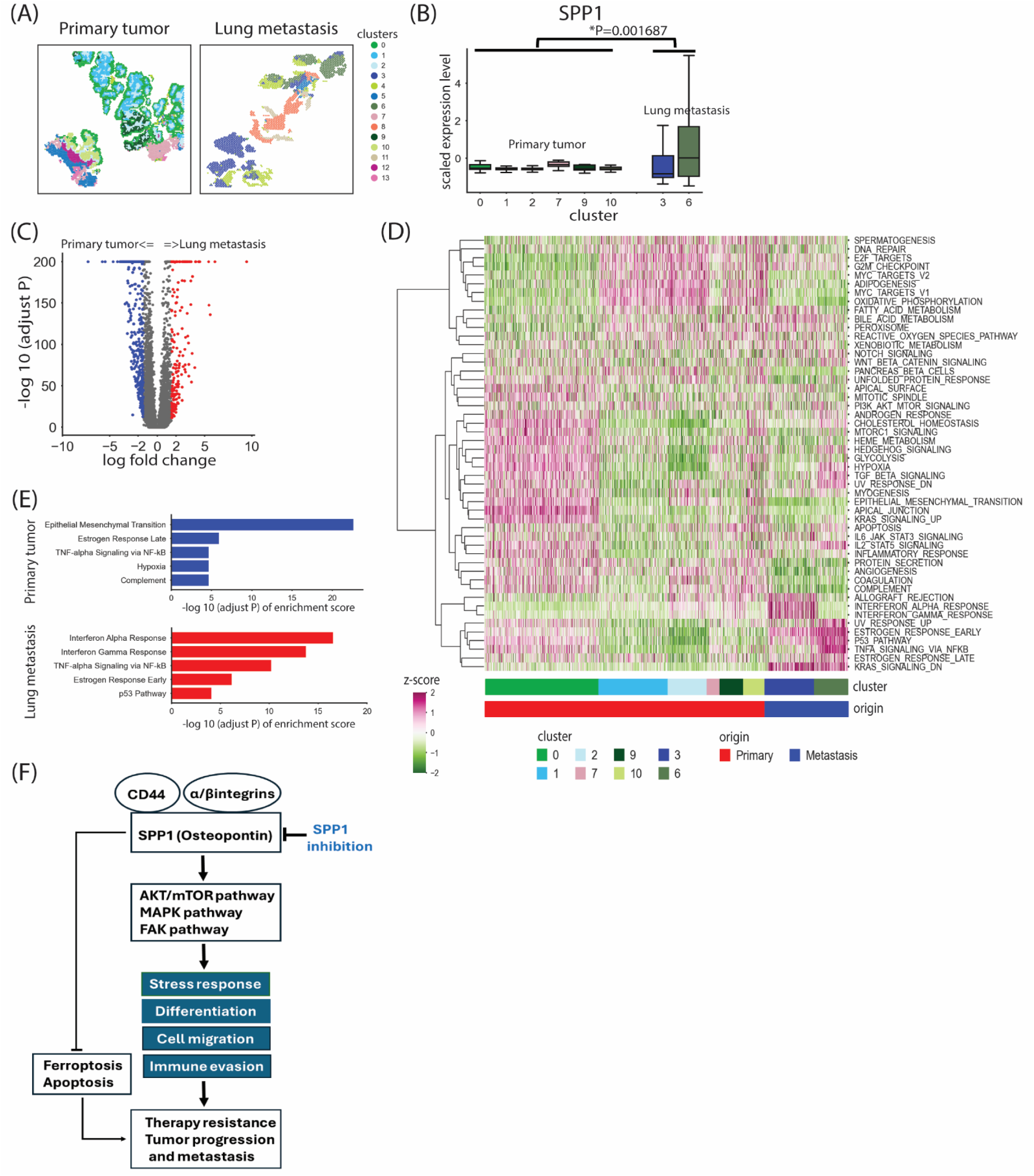
Spatial annotation and gene set enrichment analyses identify high expression of SPP1 and distinct cancer HALLMARK pathways between primary and metastatic lung tumors in HNSCC patient. **A,** All spots filtered by pathology annotation from the Visium slides analyzed by Seurat showing 14 cell clusters (0-13) in primary and metastatic lung tumors. **B,** Box plot showing scaled expression levels of SPP1 in each cell cluster (*p=0.001687, t-test). **C,** Volcano plot depicting the log fold change (logFC>1.5 and FDR <0.01) of the differentially expressed genes in primary and metastatic lung tumors. **D,** Heatmap showing HALLMARK enrichment analyses for 50 MSigDB ALLMARK gene sets profile obtained from the differentially expressed gene list of each cluster using the AUCell package. Red horizontal bar represents primary tumor with clusters (0, 1, 2, 7, 9, and 10) and blue horizontal bar represents metastatic lung tumor with clusters (3 and 6). **E,** Bar graph showing top five enriched gene signatures selected from the 50 MSigDB HALLMARK gene sets in primary and metastatic lung tumors. **F,** Proposed signaling model summarizing the oncogenic role of SPP1 in HNSCC. We hypothesized that under NRF2-stressed conditions, hyperactivated SPP1 interacts with specific class of α/β integrins and CD44 in the tumors and activate downstream signaling effectors including AKT/mTOR, MAPK and FAK proteins. Consequently, this interaction between SPP1 and its receptors in cisplatin-resistant HNSCC enhances oncogenic cellular responses and inhibits ferroptosis, ultimately contributing to therapy resistance, tumor progression, and metastasis.

## Discussion

Cisplatin-based concurrent chemo-radiotherapy represents the first-line treatment for most patients with advanced HNSCC (3). Cisplatin resistance contributes to treatment failure, development of locoregional and distant metastasis and cancer specific mortality (3). Identifying potential biomarkers that can predict the response to cisplatin is critical to decreasing the recurrence rate and fatality of HNSCC. In a prior publication, we showed that SPP1, an NRF2- target gene, is significantly upregulated *in vitro* and *in vivo* in cisplatin-resistant HNSCC tumor models (7). Several studies have shown that SPP1 contributes to the invasion and metastasis of solid tumors (23, 28–30). Using human cisplatin resistant tumor cell lines stably expressing shRNA SPP1 and patient samples, combined with high-throughput transcriptomic analysis, we systemically investigated the contribution of NRF2-upregulated SPP1 to HNSCC progression.

In this study we have clearly demonstrated that targeted suppression of SPP1 can restore the sensitivity of CDDP-resistant HNSCC tumor cells to cisplatin treatment *in vitro* and *in vivo*. We have also shown that increased sensitivity to cisplatin in SPP1 KD resistant cells is likely driven by activation of ferroptosis, a programmed cell death mechanism dependent on lipid accumulation and peroxidation via utilization of iron (31). These results further suggest that the NRF2-hyperactivated SPP1 decreases cisplatin response by suppressing ferroptosis in CDDP-resistant HNSCC cells. It is also evident in our study that targeting SPP1 effectively decreases the ability of CDDP-resistant HNSCC cells to invade and metastasize to lymph nodes and lungs *in vivo*. Collectively, these data implicate a significant and perhaps direct role for deregulated SPP1 in locoregional and metastatic progression during acquisition of cisplatin resistance in HNSCC. Furthermore, an *in vitro* competition experiment with anti-SPP1 to block interaction of SPP1 with its cell surface receptors, demonstrated that decreased tumor progression in CDDP-resistant HNSCC cells was specific to SPP1 silencing. Unlike *in vitro* data, inhibition of SPP1 with a humanized anti-osteopontin (OPN) antibody had minimal effect on tumor growth as a single agent *in vivo*. However, when combined with CDDP the antibody significantly impacted tumor growth and metastasis in CDDP-resistant HNSCC model, consistent with published data (23). Accordingly, blocking SPP1 may enhance the efficacy of cisplatin by potentially improving its ability to induce ferroptosis. Moreover, the lack of response to anti-osteopontin antibody alone we observed in our HNSCC model implies that while osteopontin (SPP1) is a tumor-promoting factor, its activity in primary tumor growth is not solely dependent on its own mechanisms but overlaps with other factors. It is also possible that the existence of two distinct families of SPP1 receptors including integrins and CD44, have the capability to trigger downstream signaling pathways independent of each other. This possibility is supported by our co-immunoprecipitation and Visium spatial transcriptomic experiments demonstrating a positive interaction between SPP1 and a class of integrins and CD44 in HNSCC tumors. Therefore, inhibition of one of the two receptor mediated pathways may not completely suppress SPP1 signaling. It is worth mentioning that we have not detected SPP1-CD44 interaction *in vitro* in the tumor cell lines. However, high resolution Visium spatial transcriptomic analysis revealed potential interaction between SPP1 and CD44 only in the metastatic lung lesions of HNSCC in patient tumors, suggesting a significant role in metastasis. This further suggests that SPP1-CD44 interaction may be context-dependent and play a more prominent role in the tumor microenvironment of metastatic HNSCC. Thus, targeting SPP1 with small molecule inhibitors, particularly those that disrupt its binding to integrin and CD44, could be a promising approach to inhibit tumor progression in NRF2-hyperactivated HNSCC.

Analysis of RNAseq data from the TCGA HNSCC cohort showed that SPP1 gene expression is an independent significant poor prognostic factor that correlates with NRF2/KEAP1 mutational status and aggressive clinical features in HNSCC. Consistent with these findings, OCSCC tumors from patients with a higher N stage (>N1) and T stage (T3 &T4) had significantly increased average expression of SPP1 and increased frequency of gross extracapsular spread. The average SPP1 expression of poorly and moderately differentiated tumors was also increased relative to well differentiated tumors. Collectively, these data provide evidence that deregulated SPP1 may be a critical NRF2-hub gene which contribute to the spread of tumor cells beyond the capsule of a lymph node into surrounding tissues. In HNSCC, extracapsular spread (ECS) is a significant prognostic factor and in most cases indicates higher risk of disease recurrence and metastasis. Importantly, the regression model revealed that NRF2 was the strongest predictor of SPP1 followed by macrophages and CAF cells. Elevated SPP1 plasma levels have been observed in head and neck cancer patients after surgery due to wound healing (32) and also after radiotherapy (33).Taken together with our data, future studies are necessary to investigate whether baseline serum and saliva levels of SPP1 can prospectively predict response to combination therapy with cisplatin plus radiation or immune checkpoint inhibitors (ICIs) in HNSCC patients with advanced disease.

To evaluate the feasibility of this clinical significance, we have been able to accurately detect substantial levels of secreted SPP1 in plasma samples obtained from small cohort of HNSCC patients (Supplementary Table S8). Using proteomic analysis, we investigated the protein changes and downstream signaling linked to SPP1in CDDP-resistant HNSCC cells. Consistent with published data (34–37), we show that silencing SPP1 inhibits activity of several critical oncogenic and metastatic downstream signaling pathways, including AKT/mTOR, PAK1 (p21-activated kinase 1) and FAK (focal adhesion kinase). FAK and PAK1 mediated- signaling pathways are deregulated in several human cancers and are linked to cell motility, tumor progression and therapeutic resistance (36,37). Of particular interest that the decrease in FAK1 and PAK1 phosphorylation levels we observed in the SPP1 knockdown HNSCC cells suggest a potential interaction and regulatory relationship between these oncogenic proteins and warrant further investigation. Spatial annotation and gene set enrichment analyses revealed differential expression of SPP1 in cell clusters with different cancer HALLMARK pathways between the primary and metastatic lung tumors in HNSCC patient. Interestingly, in primary tumor clusters, SPP1 was expressed at low levels and notably enrichment of EMT-related genes. However, in metastatic lung tumor clusters, SPP1 is highly enriched, along with interferon response gene signatures. Mechanistically, these findings implicate that SPP1 contributes to tumor progression, initially by driving tumor growth through regulation of EMT, and then by influencing tumor metastasis through interactions within the tumor microenvironment (TME). Consistent with our data, recent study has demonstrated that SPP1 promotes cancer cell growth and resistance to chemoradiotherapy through induction of EMT (38). These functions are mediated by activation signals from the PI3K/Akt and MAPK pathways induced by binding of SPP1 to integrin αvβ3 and CD44 (38). It is reasonable to postulate that co-expression of SPP1 and interferon genes reported in in our study, participates in augmenting inflammatory responses that favor immunosuppression, and therefore facilitating tumor metastasis in NRF2-activated HNSCC.

SPP1 plays a significant role in metastasis by influencing macrophage behavior and function within the tumor microenvironment (39,40). Specifically, SPP1+ macrophages, have been linked to different aspects of immune suppression, tumor progression, including early lymph node metastasis and enhanced angiogenesis (39,40). Taken together, future studies are needed to systemically investigate whether SPP1 expression patterns promote immune tolerance and suppress immune cell activation in NRF2-hyperactivated head and neck tumors. In summary, our data demonstrated that specific targeting of SPP1 inhibits tumor progression in Nrf2 hyperactivated cisplatin resistant HNSCC. Furthermore, our study suggests that SPP1 may represent a diagnostic and prognostic biomarker, as well as a potential therapeutic target in HNSCC. SPP1 is frequently overexpressed in multiple compartments of the TME, such as in tumor associated macrophages (TAMs) or cancer associated fibroblasts. Therefore, additional experiments are needed to disentangle the contribution of SPP1 expressed in the TAM and CAF cells to development of HNSCC tumor metastasis in the presence and absence of NRF2 hyperactivation. Importantly, alternative protein translation produces an intracellular form of SPP1 and a secreted form that can bind ubiquitous integrin receptors or CD44 receptors on tumors and other cell types to activate signaling cascades. Thus, functional experiments are also required to clarify differences in the impact of secreted versus intracellular SPP1 in NRF2-hyperactivated HNSCC tumor cells.

## Abbreviations

HNSCC: head and neck squamous cell carcinoma
CDDP: cisplatin
KEAP1: Kelch-like ECH-associated protein 1
NRF2: nuclear factor erythroid 2–related factor 2
CUL3: Cullin 3
OCSCC: oral cavity squamous cell carcinoma
SPP1: secreted phosphoprotein 1 OPN (osteopontin)
GPX4: (glutathione peroxidase 4)
ACSL4: acyl-CoA synthetase long chain family member 4.

---

**Supplementary Figure S1.**
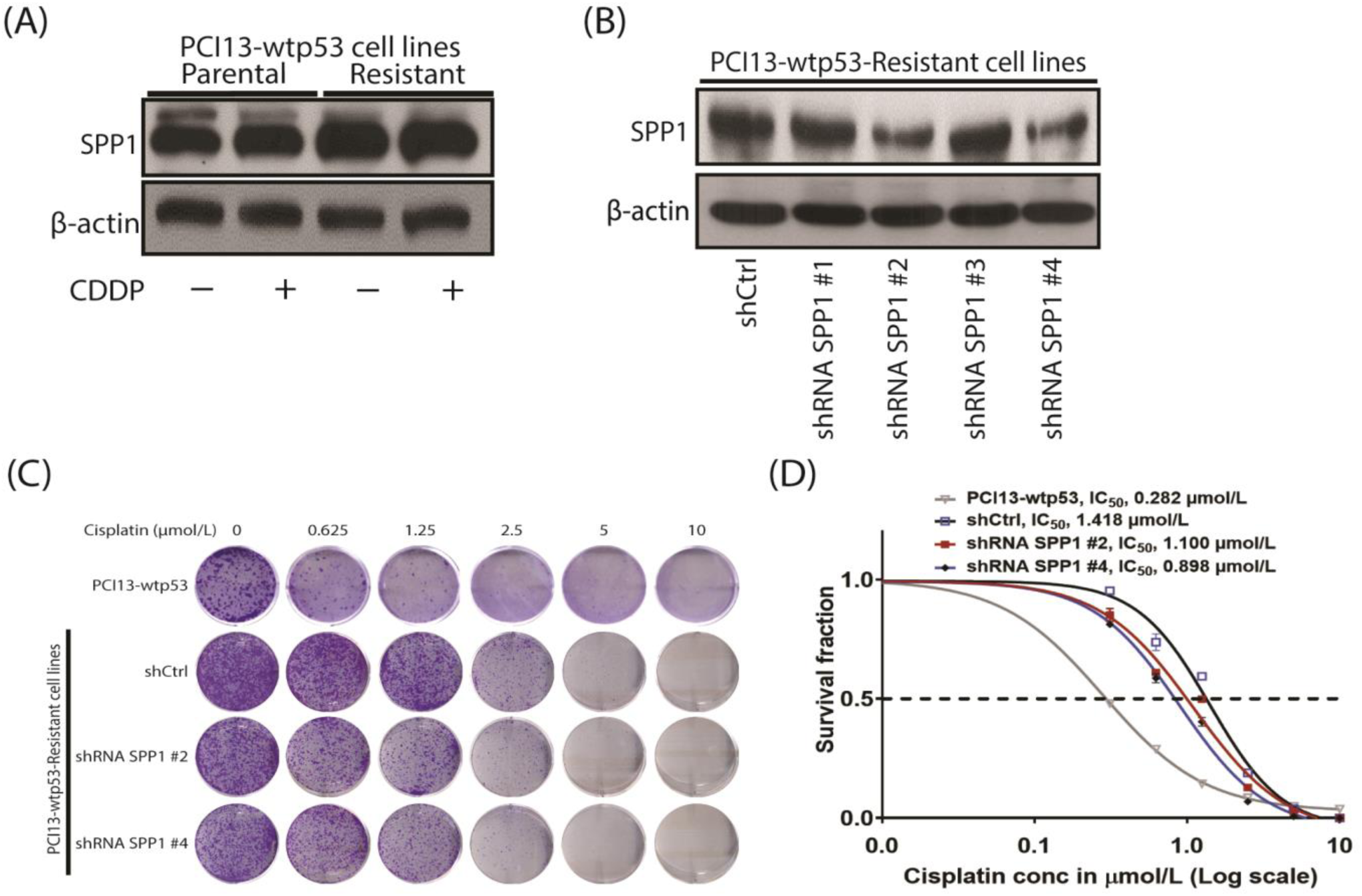
Downregulation of SPP1 enhances cisplatin sensitivity *in vitro* in cisplatin resistant HNSCC. **A,** Western blot demonstrates high expression of SPP1 during acquired CDDP resistance in PCI13-wtp53 HNSCC cells compared to PCI13-wtp53 CDDP sensitive parental cells. **B**, Western blot verifies downregulation efficiency of SPP1 in stable CDDP resistant cells (PCI13-wtp53-resistant). **C and D,** Representative images of clonogenic survival and curves demonstrating increased sensitivity to CDDP in PCI13-wtp53 cells stably expressing shRNA SPP1 after treatment with the indicated doses of cisplatin.

**Supplementary Figure S2A.**
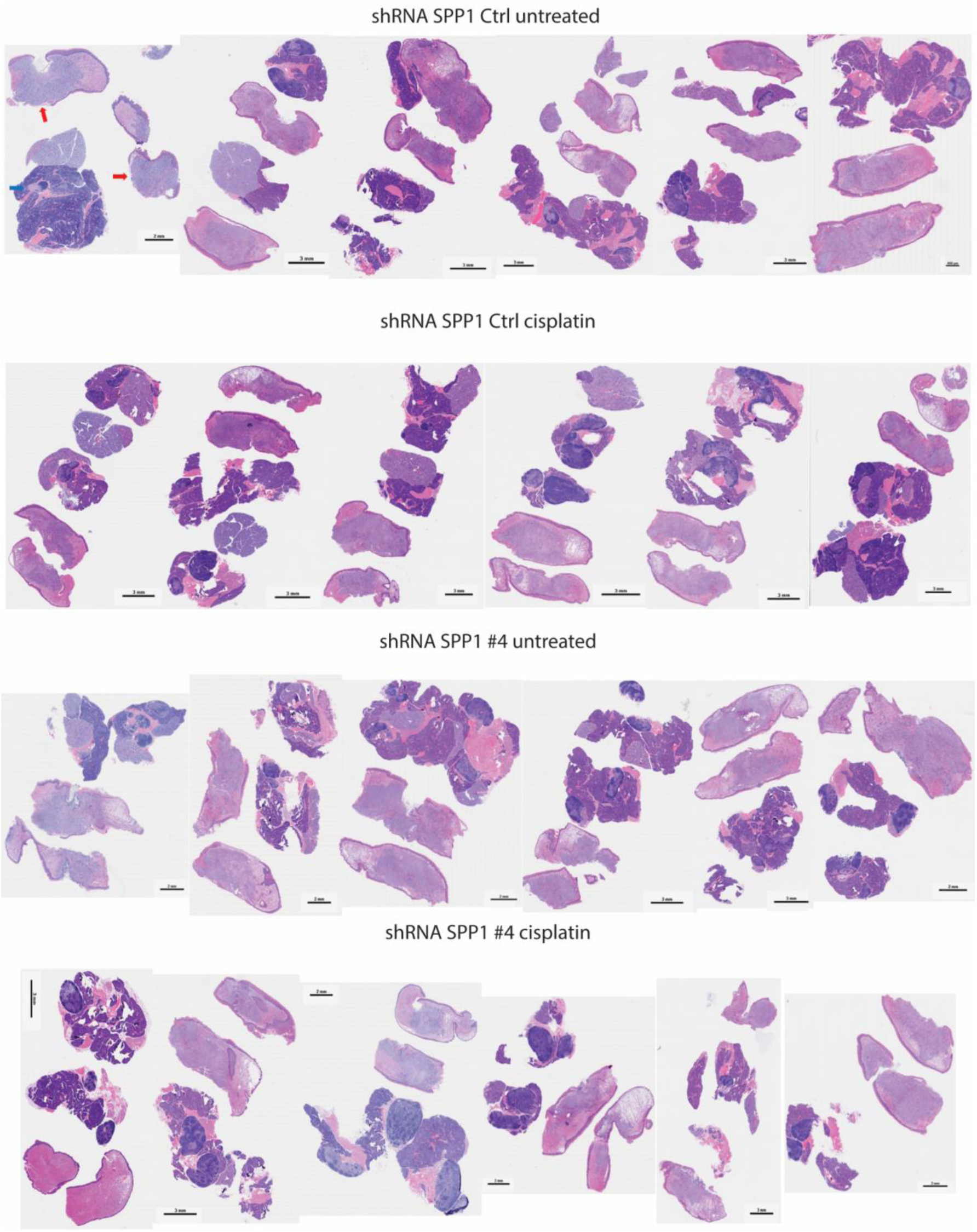
Tongue and lymph node harvested from the mice injected with HN30-R8 cells stably expressing shRNA Ctrl and shRNA SPP1 following cisplatin treatment. Red arrows indicate tongue tumor, and blue arrows indicate cervical lymph node.

**Supplementary Figure S2B.**
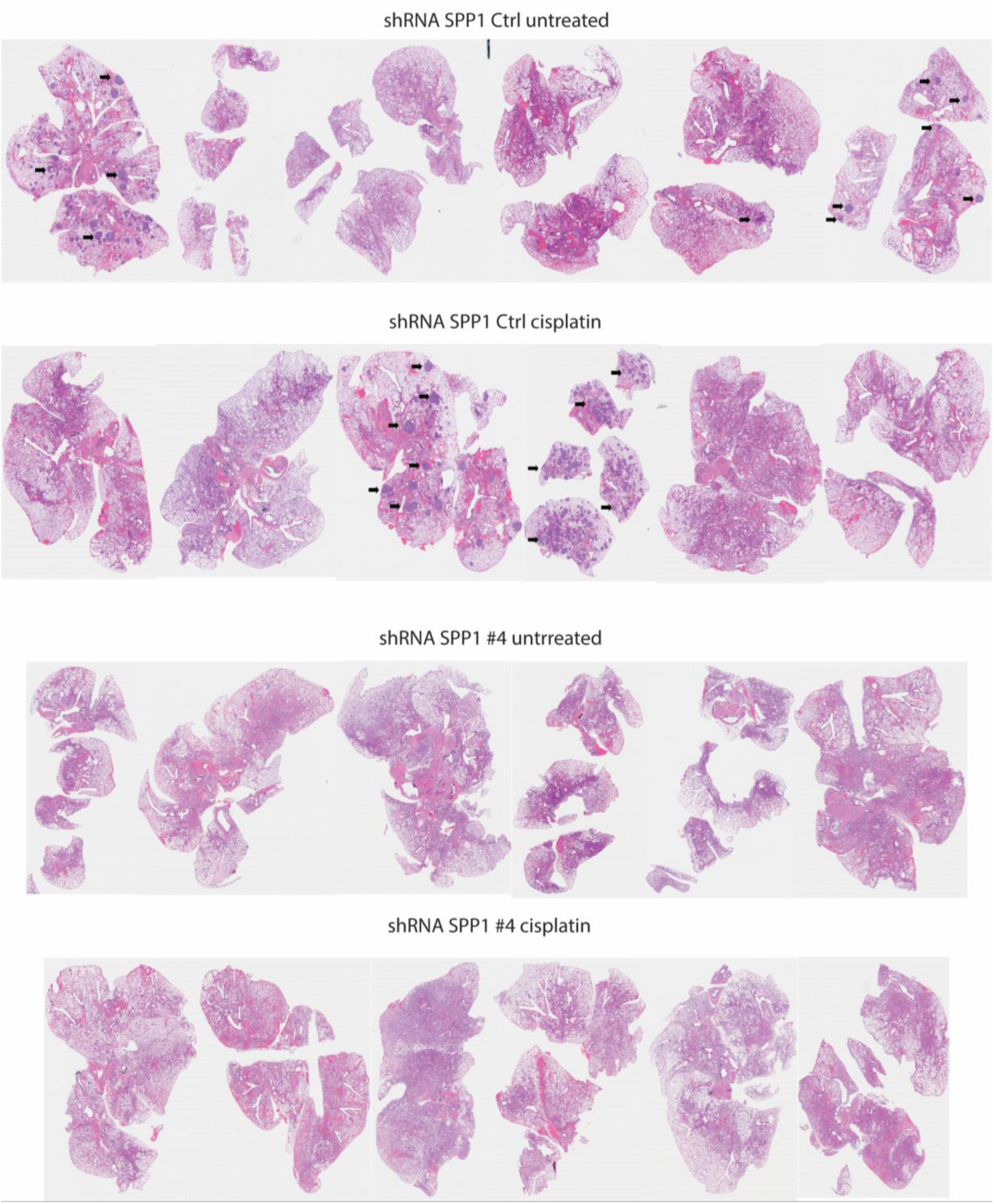
Lungs harvested from the mice injected with HN30-R8 cells stably expressing shRNA Ctrl and shRNA SPP1 following cisplatin treatment. Black arrows indicate metastatic lung tumor nodules.

**Supplementary Figure S3A.**
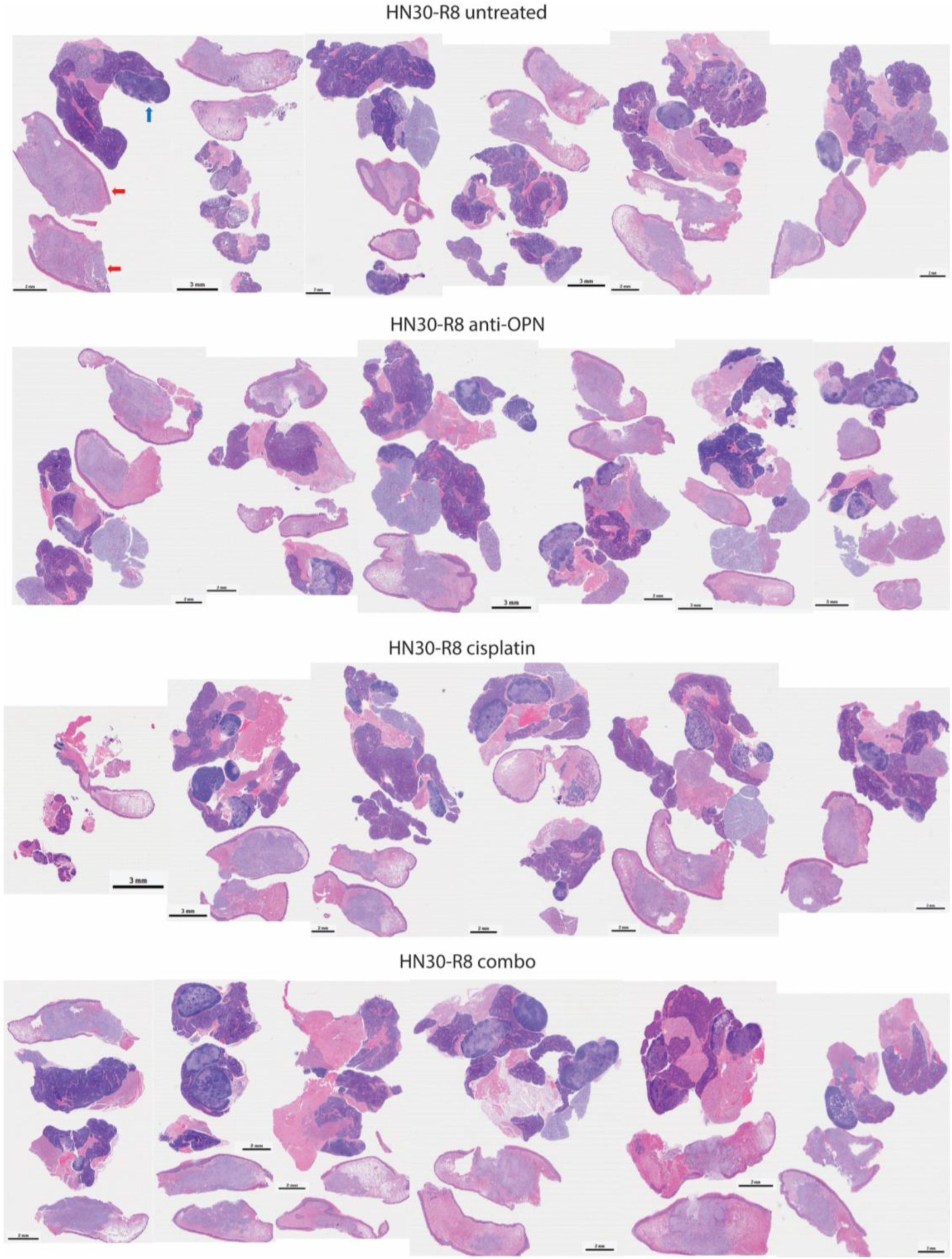
Tongue and lymph node harvested from the mice injected with HN30-R8 cells following treatment with cisplatin in combination with anti-SPP1 (anti-osteopontin). Red arrows indicate tongue tumor, and blue arrows indicate cervical lymph node.

**Supplementary Figure S3B.**
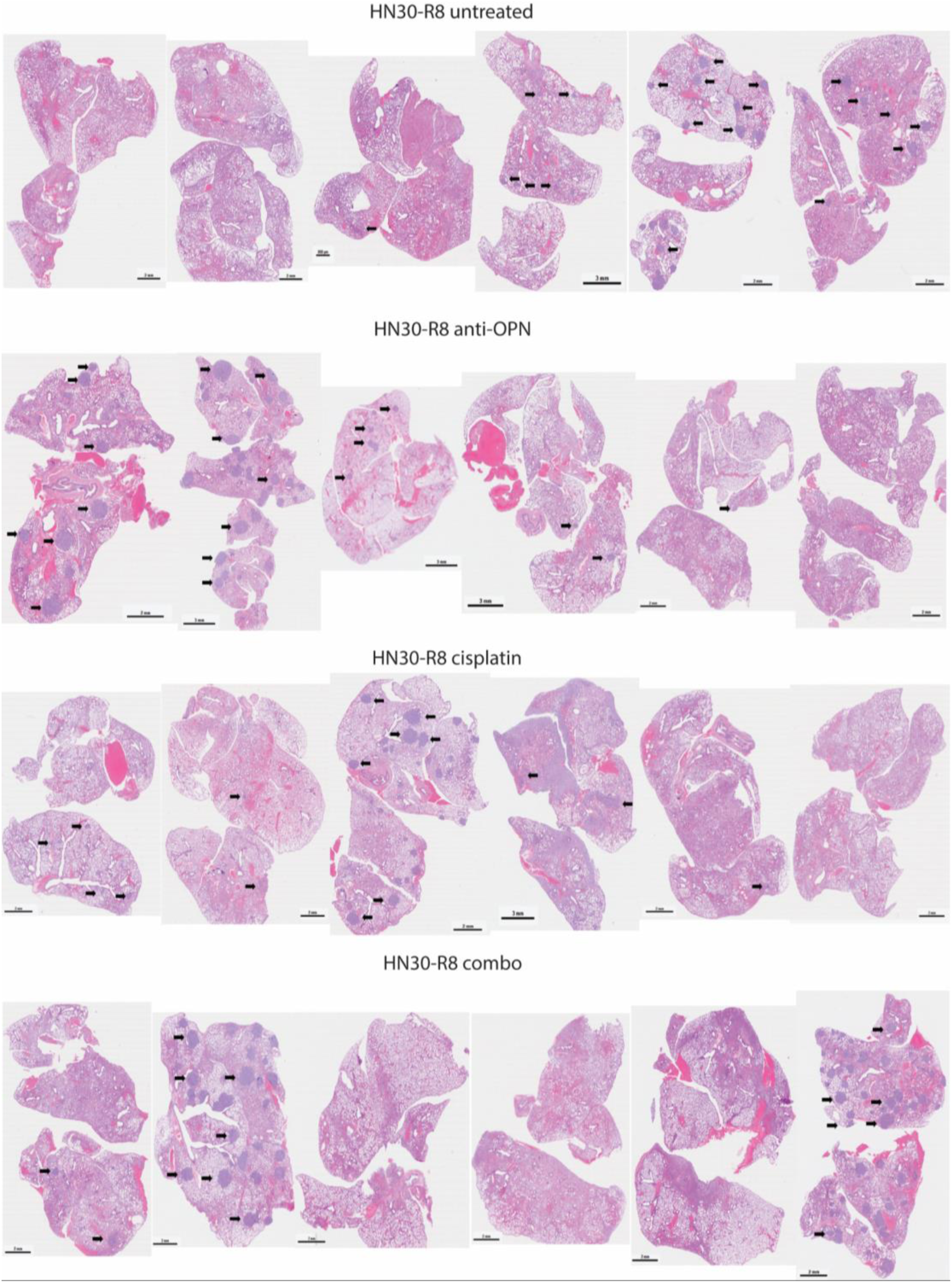
Lungs harvested from the mice injected with HN30-R8 cells following treatment with cisplatin in combination with anti-SPP1 (anti-osteopontin). Black arrows indicate metastatic lung tumor nodules.

**Supplementary Figure S4.**
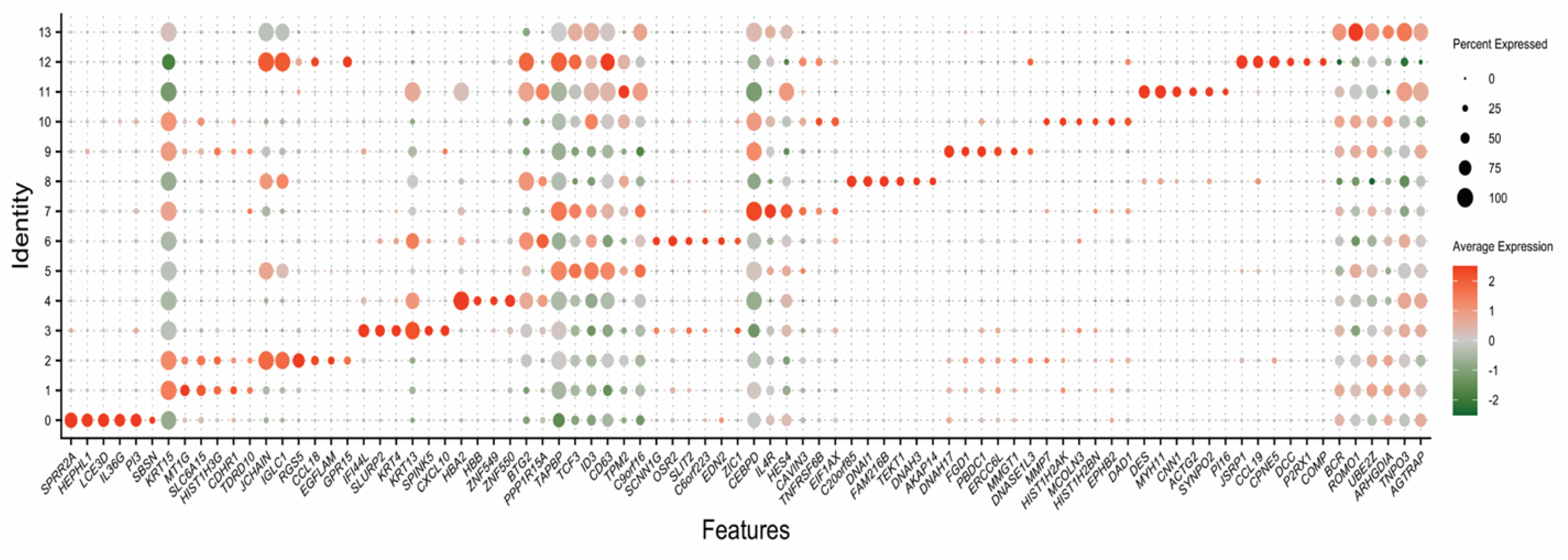
Bubble plot annotating identified cell types and clusters expressing the top 6 significantly expressed genes in each cluster.

## Notes

**Financial Support:** This work was supported by the National Institute of Health/NCI U54CA27432 (A.A. Osman and J.N. Myers) and Cellular Imaging Core, Bioinformatics Shared Resource, and the Reverse Phase Protein Array (RPPA) Core, which are supported by the National Institutes of Health through MD Anderson’s Cancer Center Support Grant (P30CA016672).

**Conflict of Interest Disclosure:** The authors declare no conflict of interest.

### Competing Interest Statement

The authors have declared no competing interest.

