## Supplementary Materials and Methods for "Targeted suppression of SPP1 inhibits tumor invasion and metastasis in NRF2 hyperactivated cisplatin resistant HNSCC"

Kawabe M et al.

### Supplementary Materials and Methods

#### Generation of SPP1 knockdown stable cell lines.

The lentiviral shRNA plasmids, pGIPZ-NT (Cat# RHS4348), pGIPZ-SPP1#1 (Clone # V2LHS_303525), pGIPZ-SPP1#2 (Clone # V2LHS_303526), pGIPZ-SPP1#3( Clone # V2LHS_303528), pGIPZ-SPP1#4 (V2LHS_111534) were all obtained from Dharmacon. HN30-R8 cells stably expressing the lentiviral non-targeting (NT)control and the pGIPZ-SPP1 plasmids were generated as previously described (7). Briefly, virus containing supernatants were collected 48 hours and 72 hours after co-transfection of the pGIPZ-NT, pGIPZ-SPP1 plasmid and the lentiviral packaging vectors pCMV-dR8.2 dvpr and pMD2.G into HEK293-FT cells (using Lipofectamine 2000). HN30- R8 target cells were infected with the virus in the presence of polybrene (polybrene (5 µg/mL) and cells were selected first with puromycin ((1.0 µg/mL) followed by sorting of GFP-positive cells. SPP1 knockdown in the stable cell lines were confirmed by Western blotting. The clones showed substantial degree of SPP1 knockdown (Clone #2 and #4) were established and used for further analysis.

#### List of Antibodies used for western blotting

Antibodies used for Western blotting and immunoprecipitation were SPP1 (#22952-1-AP, Proteintech); NRF2 (ab137550, Abcam); KEAP1 (#ab119403, Abcam); PARP-1 (#9542), FAK1 (#13009), phospho-FAK1 (#8556), CD44 (#3570); all from Cell Signaling Technology; GPX4 (E-12), (#SC-166570), ACSL4 (#SC-271800), Integrin α4/ITGA4/CD49d (B-2) (#SC-52593),

Integrin αV/ITGAV/CD51 (P2W7) (#SC-9969), Integrin β1/ITGB1 (A-4) (#SC-374429), Integrin β 6 (C-19) (SC-6632), and Integrin β3/ITGB3/CD61 (D-11) (#SC-365679); all from Santa Cruz Biotechnology; β-actin (#A2228; Millipore Sigma).

**Reverse phase protein array (RPPA)**

Samples were prepared as described previously (18). Briefly, HN30-R8 cells stably expressing lentiviral vector control (shCtrl) and SPP1 knockdown derivative (SPP1 shRNA #4) were seeded in 10-cm dishes and harvested at 24 hours with RIPA lysis buffer containing protease and phosphatase inhibitors as indicated. Protein lysates were then collected, vortexed, centrifuged and total protein resulting from supernatant for each sample was quantitated using a BCA kit (Pierce Biotechnology Inc., Rockford, IL). RPPA lysates were prepared in biological triplicate RPPA was performed by the RPPA core facility at the University of Texas MD Anderson Cancer Center. Protein expression data were generated by RPPA for 77 protein analytes. RPPA slides were quantified using ArrayPro (Media Cybernetics) to generate signal intensities that were further processed by SuperCurve to estimate relative protein levels (in log2 scale). RPPA samples quality was monitored by a QC classifier and only the slides whose QC scores were above 0.8 (on a 0–1 scale) were used for further analysis. Differences in group mean RPPA analytes were identified with independent two-sample (two-sided) t-tests using JMP13 software, adjusting for multiple comparisons with the Benjamini-Hochberg procedure ($q \leq 0.1$). Volcano plots of significance verses differences in RPPA level were generated with GraphPad Prism software, and two-way Ward's hierarchical clustering for select analytes with the corresponding heatmap was generated in JMP. Significant differentially expressed proteins and phosphoproteins among the groups were subjected to Gene Ontology (GO) enrichment analysis to identify cell function.
