## Supplementary Table S5 for "Targeted suppression of SPP1 inhibits tumor invasion and metastasis in NRF2 hyperactivated cisplatin resistant HNSCC"

| Primary tumor | | cluster | | | | | | | | | | | | |
|---|---|---|---|---|---|---|---|---|---|---|---|---|---|---|
| | | 0 | 1 | 2 | 4 | 5 | 6 | 7 | 8 | 9 | 10 | 11 | 12 | 13 |
| Pathology Annotations | Bacterial colonies | 1 | 0 | 0 | 1 | 0 | 0 | 0 | 0 | 0 | 0 | 0 | 0 | 0 |
| | Erythrocytes | 73 | 5 | 1 | 9 | 4 | 0 | 1 | 0 | 0 | 0 | 0 | 0 | 10 |
| | Hemorrhage | 49 | 2 | 0 | 53 | 4 | 0 | 3 | 0 | 0 | 0 | 0 | 0 | 4 |
| | Lamina propria | 0 | 0 | 0 | 0 | 36 | 0 | 2 | 0 | 0 | 0 | 0 | 5 | 13 |
| | Lymphoplasmacytic infiltrate | 4 | 3 | 218 | 1 | 54 | 0 | 29 | 1 | 0 | 5 | 0 | 132 | 0 |
| | Squamous cell carcinoma | 1415 | 864 | 488 | 4 | 39 | 6 | 163 | 0 | 291 | 269 | 1 | 0 | 15 |
| | Unclassified | 4 | 0 | 1 | 9 | 241 | 0 | 216 | 0 | 1 | 0 | 0 | 0 | 65 |
| | Vessel | 1 | 18 | 63 | 0 | 16 | 0 | 1 | 1 | 1 | 0 | 0 | 6 | 0 |

| Lung metastasis | | cluster | | | | | | | |
|---|---|---|---|---|---|---|---|---|---|
| | | 0 | 2 | 3 | 4 | 5 | 6 | 8 | 11 |
| Pathology Annotations | Bronchial epithelium | 0 | 0 | 0 | 0 | 1 | 0 | 162 | 1 |
| | Erythrocytes | 0 | 0 | 2 | 35 | 0 | 15 | 4 | 0 |
| | Fibroconnective tissue | 0 | 0 | 2 | 39 | 49 | 0 | 210 | 141 |
| | Hemorrhage | 1 | 0 | 6 | 294 | 1 | 41 | 1 | 0 |
| | Lymphoplasmacytic infiltrate | 0 | 0 | 0 | 0 | 0 | 0 | 22 | 0 |
| | Mucin | 0 | 0 | 0 | 3 | 0 | 0 | 0 | 0 |
| | Necrosis | 0 | 0 | 0 | 0 | 0 | 3 | 0 | 0 |
| | Squamous cell carcinoma | 1 | 1 | 619 | 31 | 12 | 370 | 1 | 0 |
| | Vessel | 0 | 1 | 2 | 0 | 3 | 6 | 12 | 1 |

**Supplementary Table S5. Pathological annotation of the clusters in primary and metastatic lung tumors analyzed by Seurat.** The tabulated annotation shows the population of cell types in each cluster present in the primary and metastatic lung tumors. Clusters with higher number of squamous cell carcinoma type were selected for analysis.
