## Supplementary Table S8 for "Targeted suppression of SPP1 inhibits tumor invasion and metastasis in NRF2 hyperactivated cisplatin resistant HNSCC"

| n=28 | SPP1 (ng/ml) | *P-value* (t-test) |
|---|---|---|
| REC/DM | 129.1 | 0.38 |
| none | 110.9 | |
| HP | 165.9 | |
| LA | 98.2 | |
| OC | 115.8 | |
| OP (p16+) | 97.9 | |
| T1-2 | 113.3 | 0.43 |
| T3-4 | 120.0 | |
| N0 | 107.5 | 0.45 |
| N>0 | 121.8 | |
| smoker | 122.7 | 0.23 |
| non-smoker | 98.0 | |

**Supplementary Table S8. ELISA analysis of SPP1 (osteopontin) in plasma obtained from HNSCC patients.** Plasma samples were processed and subjected to ELISA analysis as described in the Methods section according to manufacturer's procedures following dilution (1:25) in dilution buffer. SPP1 levels were high in all patients and trended toward correlation with smoking status but this did not reach statistical significance. REC/DM= subsequently developed locoregional recurrence and/or distant metastasis; HP= hypopharynx; LA= hypopharynx; OC= oral cavity; OP= oropharynx. P-values denote comparison between individual categories using a 2-sided t-test.
